# A High-Resolution Landscape of Mutations in the *BCL6* Super-Enhancer in Normal Human B-Cells

**DOI:** 10.1101/783118

**Authors:** Jiang-Cheng Shen, Ashwini S. Kamath-Loeb, Brendan F. Kohrn, Keith R. Loeb, Bradley D. Preston, Lawrence A. Loeb

## Abstract

The super-enhancers (SE) of lineage-specific genes in B-cells are off-target sites of somatic hypermutation. However, the inability to detect sufficient numbers of mutations in normal human B-cells has precluded the generation of a high-resolution mutational landscape of SEs. Here, we captured and sequenced 12 B-cell SEs at single-nucleotide resolution from ten healthy individuals across diverse ethnicities. We detected a total of ∼9000 subclonal mutations (allele frequencies <0.1%); of these, ∼8000 are present in the *BCL6* SE alone. Within the *BCL6* SE, we identified three regions of clustered mutations where the mutation frequency is ∼7X10^-4^. Mutational spectra show a predominance of C>T/G>A and A>G/T>C substitutions, consistent with the activities of activation-induced-cytidine deaminase (AID) and the A-T mutator, DNA Polymerase η, respectively, in mutagenesis in normal B-cells. Analyses of mutational signatures further corroborate the participation of these factors in this process. Single base substitution signature SBS85, SBS37, and SBS39 were found in the *BCL6* SE. While SBS85 is a denoted signature of AID in lymphoid cells, the etiologies of SBS37 and SBS39 are still unknown. Our analysis suggests the contribution of error-prone DNA polymerases to the latter signatures. The high-resolution mutation landscape has enabled accurate profiling of subclonal mutations in B-cell SEs in normal individuals. By virtue of the fact that subclonal SE mutations are clonally expanded in B-cell lymphomas, our studies also offer the potential for early detection of neoplastic alterations.

**Significance:** We used Duplex Sequencing to detect low-frequency mutations in the *BCL6* super-enhancer locus in normal human B-cells. The landscape of pre-existing mutations is remarkably conserved across different ethnicities and reveals clustered mutational hotspots that correlate with reported sites of clonal mutations and translocation breakpoints in human B-cell lymphomas. This high-resolution genomic landscape revealed by Duplex Sequencing offers accurate and thorough profiling of low frequency, pre-existing mutations in normal individuals, and the potential for early detection of neoplastic alterations.

## Introduction

In response to antigen stimulation, naïve B-cells congregate in germinal centers where they undergo multiple rounds of cell division and antigenic selection ultimately maturing into antibody-producing plasma cells and memory B-cells (1–3). In the germinal center somatic hypermutation (SHM) occurs targeting the variable domain of the immunoglobulin receptor resulting in either non-functional, low or high affinity antibodies. This process involves the creation of multiple single-base substitutions in immunoglobulin genes by the mutagenic SHM machinery (4). SHM includes two stages of mutagenesis – cytidine deamination by activation-induced cytidine deaminase (AID) to generate a G:U mispair and subsequent error-prone repair synthesis by an error-prone DNA polymerase. Although SHM is a tightly regulated process, it can act aberrantly, even in normal cells, to somatically mutate other (off-target) sites. Aberrant somatic hypermutation (aSHM) sites are frequently found in non-coding DNA regions containing cis regulatory elements such as intragenic super-enhancers (SE) – binding sites for multiple factors that increase transcription (5–7). These regions are proximal to the transcription start site (TSS) of actively transcribed genes and are in an ‘open’ chromatin conformation (p300 and H3K27ac positive).

Aberrant somatic hypermutation targets were delineated by low resolution single cell-based PCR sequencing of sites with clustered mutations (8). The first of these targets to be identified was the promoter/super enhancer region of the *BCL6* gene (8, 9). The *BCL6* gene encodes a transcription corepressor protein, whose expression is tightly coordinated with entry and exit from germinal centers (10, 11). Germinal center B-cells have high levels of the BCL6 protein where it regulates the expression of many genes involved in B-cell differentiation. On the other hand, antibody-secreting plasma cells and memory B-cells that exit the germinal center turn off *BCL6* expression to facilitate the switch between differentiation and stable cell-state maintenance. It was hypothesized that mutations within the *BCL6* SE dysregulate *BCL6* expression to promote lymphomagenesis (12, 13). Indeed, Sanger sequencing of PCR amplicons from diffuse large B-cell lymphomas has identified clonal mutations in the *BCL6* SE locus (12). Further, reporter assays in cultured cells showed that some of these mutations interfere with the binding of transcriptional repressor factors to upregulate *BCL6* expression (14). More recently, Next Generation Whole Genome Sequencing studies of diffuse large B-cell lymphomas (DLBCL) reported the presence of mutation clusters within the *BCL6* SE locus (7), however, only about 30 clonal mutations were observed in a total of ten lymphoma samples, which is inadequate for characterizing the types and distributions of mutations.

A high-resolution landscape of SE mutations in B-cells from healthy individuals will aid to unravel the process of aSHM and enable early detection of pre-existing SE mutations that might signal lymphomagenesis. We reasoned that DNA mutated by aSHM is preserved in circulating memory B-cells such that deep sequencing of purified B-cells would reveal details of the mutational landscape. We employed Duplex Sequencing (DS) (15, 16) for this purpose. DS provides a mutational landscape of genomic DNA at single-nucleotide resolution to reveal mutational patterns and potential underlying mechanisms. The accuracy of DS stems from copying both strands of single DNA molecules; mutations are defined based on complementarity and presence in both strands at the same position. As a result, DS is approximately a thousand-fold more accurate than routine next-generation sequencing and allows the identification of rare mutations – those present at frequencies as low as one base-substitution in 10^7^ nucleotides sequenced.

Using targeted capture, we purified and sequenced twelve SE loci in genomic DNA samples of human CD19+ B-cells isolated from ten healthy individuals of diverse ethnic backgrounds. The mutational landscape shows clustered mutations, most elevated in the SE of *BCL6*, and to a lesser extent in the SEs of *PAX5* and *CD83.* A total of ∼9,000 base substitutions were detected in the ten samples; ∼8,000 of these were found in the *BCL6* SE. The mutational profiles of the *BCL6* SE are remarkably similar in individuals across various ethnic groups, suggesting that they are not the result of random stochastic events but instead may function in normal developmental processes. Mutational spectra and signatures at the *BCL6* SE locus further suggest that mutations result from the activities of AID and of the error-prone DNA polymerase, Pol η.

## Results

### High frequency of mutations in the *BCL6* super-enhancer

We sequenced 12 aberrant somatic hypermutation targets (Table S1) in CD19+ B-cells from ten healthy blood donors of different ethnicities (Table S2). The target gene segments are located ∼1 kb 3’ of the transcription start site (TSS), and are reported super-enhancers of the associated genes (5–7). As a control, we also captured and sequenced a SHM target locus frequently found in memory B-cells – the variable region of the immunoglobulin gene, *IgHV3-23* (17, 18). Of the 12 sequenced SE loci, we found that *BCL6* is the most frequently mutated. We observed more than 8,000 subclonal mutations (allele fraction < 0.5%) in total in 10 individuals to yield an average mutation frequency (MF = mutations/total nucleotides sequenced) of 2.2 x 10^-4^ (Fig. 1 and Table S3), comparable to that observed at the SHM target, *IgHV3-23* (MF = 1.3 x 10^-4^, Fig. 1 and Table S3). Mutant allele fractions (MAF = number of mutations at each nucleotide position/total alleles sequenced at each nucleotide position) ranging from ∼0.002% to ∼0.5% were detected in the *BCL6* SE (Figs. 2a and 2b). Since circulating memory B-cells in peripheral blood comprise between 5-20% of the B-cell population (19), we estimate that there are between 0.8-3.2 mutations within the 1 kb *BCL6* SE region in each circulating memory B-cell, assuming that other circulating B-cells do not contain frequent mutations in *BCL6*.

**Figure 1.**
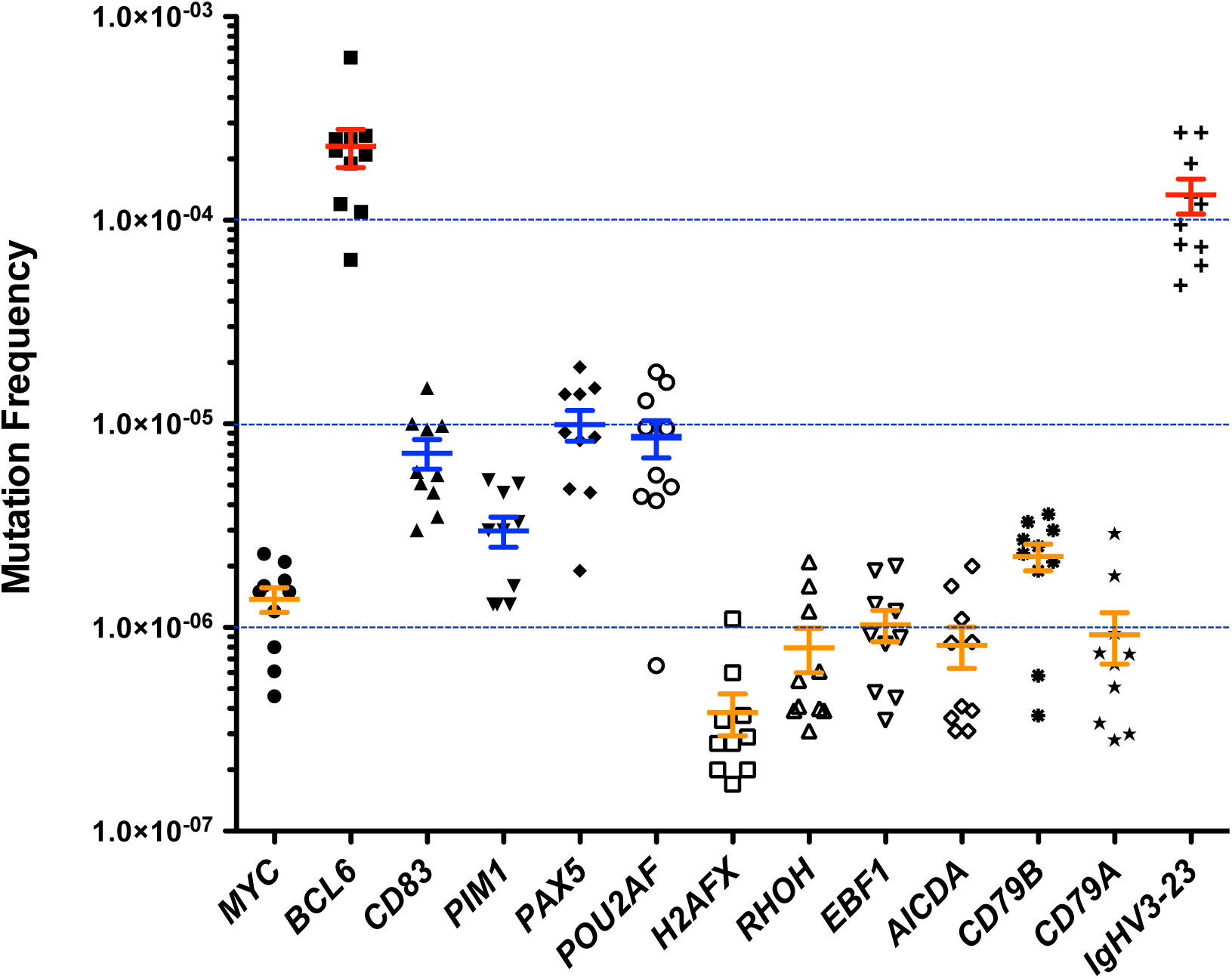
Mutation frequencies in various TSS (transcription start site)-SE gene loci and in the immunoglobulin gene *IgHV3-23* in B-cells from ten healthy individuals. Twelve aSHM targets as well as the immunoglobulin gene *IgHV3-23* were captured from CD19+ B-cells purified from ten independent blood donors and sequenced by Duplex Sequencing. Mutation frequencies – total number of mutations divided by total number of nucleotides sequenced, were calculated for each sample and plotted as shown. Note that the mutation frequency at the *BCL6* SE (2.2 x 10^-4^) is as high as that of *IgHV3-23* (1.3 x 10^-4^). Data are presented as mean ± SEM (N=10). SEs with mutation frequencies greater than 1.0 x 10^-4^, between 2.5 x 10^-6^ and 1.0 x 10^-4^, and less than 2.5 x 10^-6^ are highlighted by red, orange, and blue marks, respectively.

**Figure 2.**
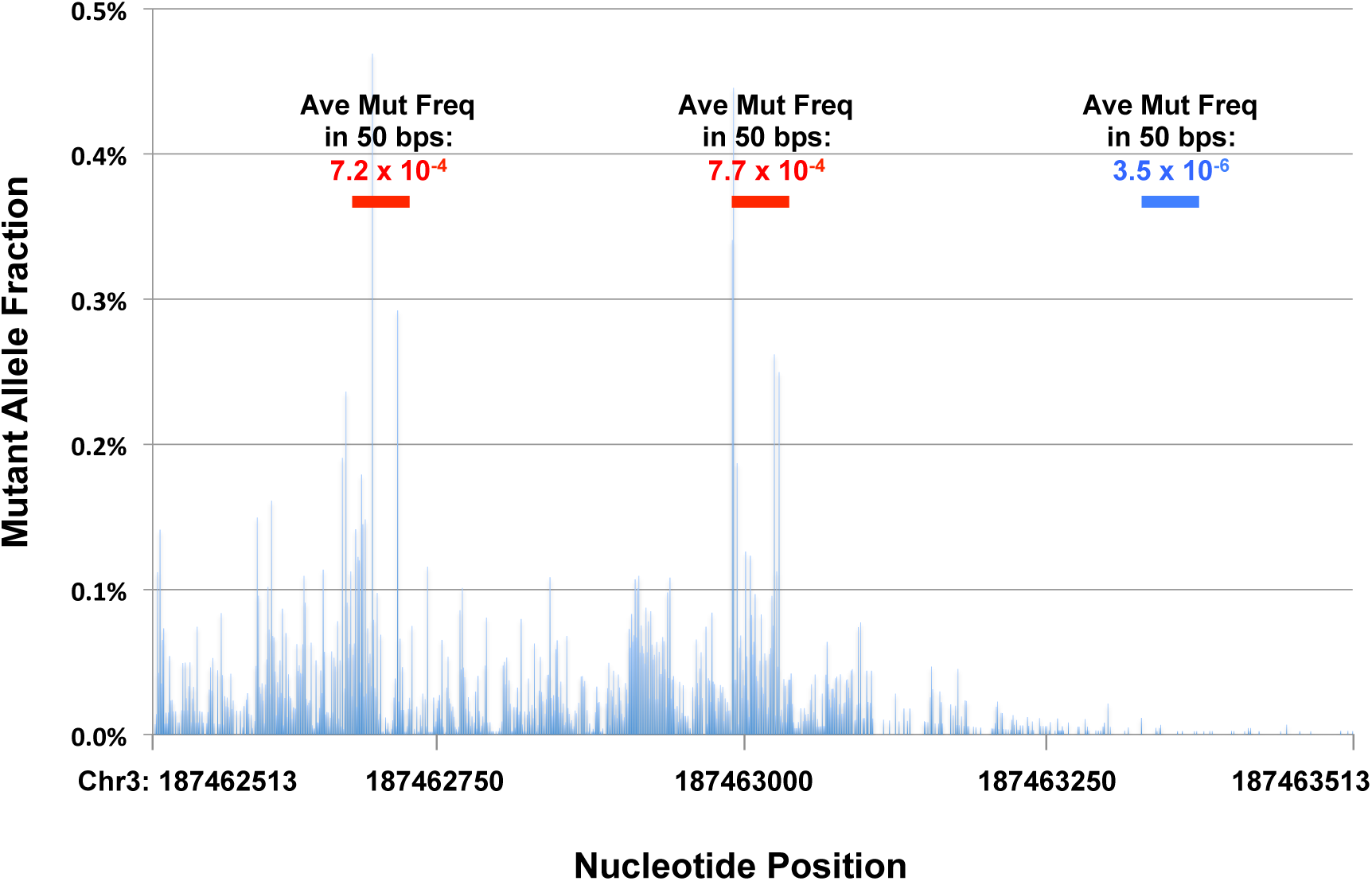
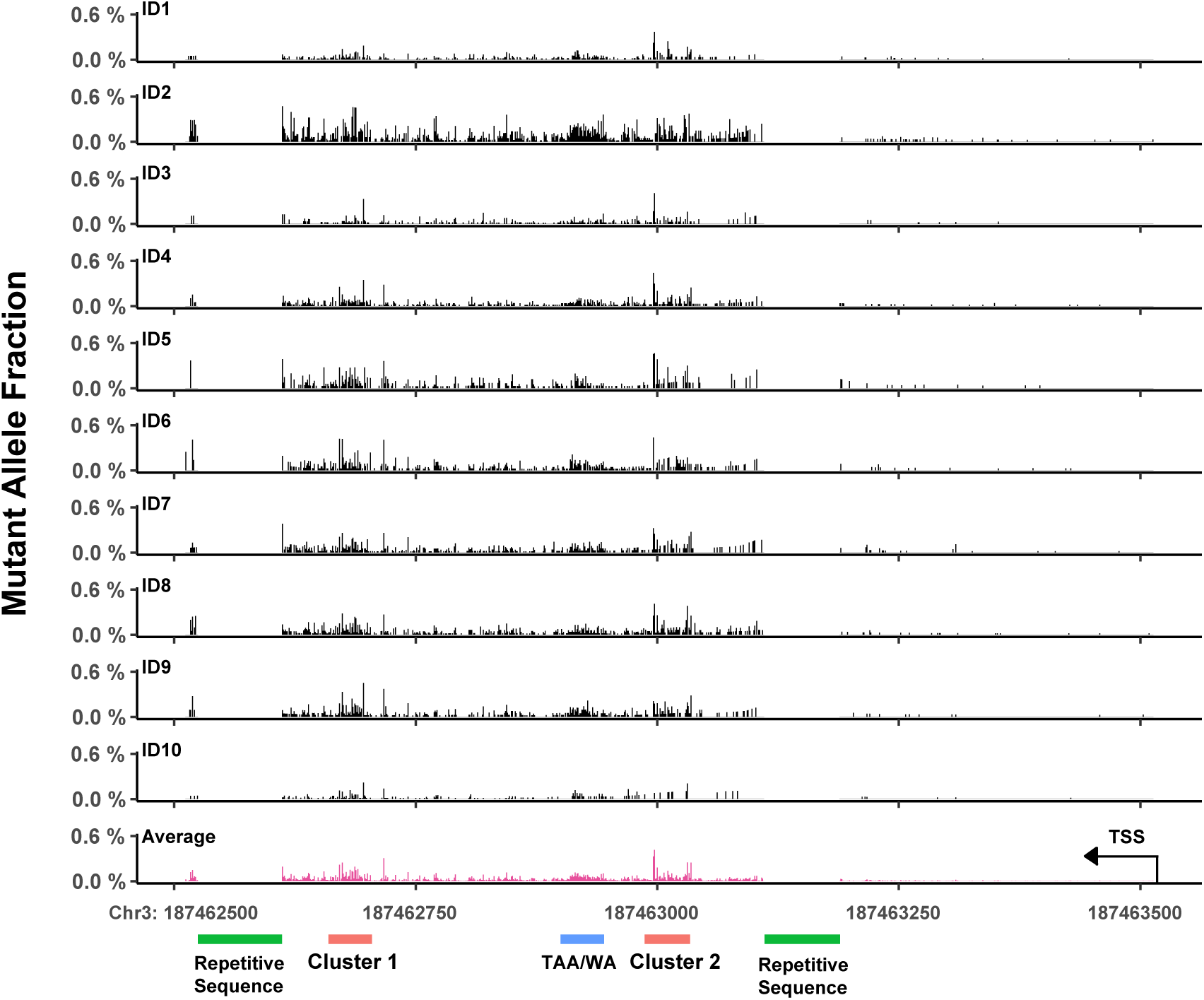
Landscape of aberrant somatic hypermutation at the *BCL6* super-enhancer locus. a) Average mutant allele fractions (MAF; number of mutations at a given nucleotide position/sequence read depth at that position) at each nucleotide position within the 1,001 bp region of the *BCL6* SE. Average mutant allele fractions in the two 50-bp hotspot clusters and in a 50-bp cold spot are indicated. b) Mutational profiles of the *BCL6* SE locus are conserved in individuals of different ethnicities. Most of the mutations are present in the 600-bp segment at the 5’-end of the genomic DNA sequence, i.e. between positions +400 and +1,000 from the transcription start site (TSS). Three mutational clusters are apparent. Whereas mutations in clusters 1 and 2 have higher mutant allele fractions, those in the TAA/WA sequence have lower fractions but higher density.

**Table 1.**
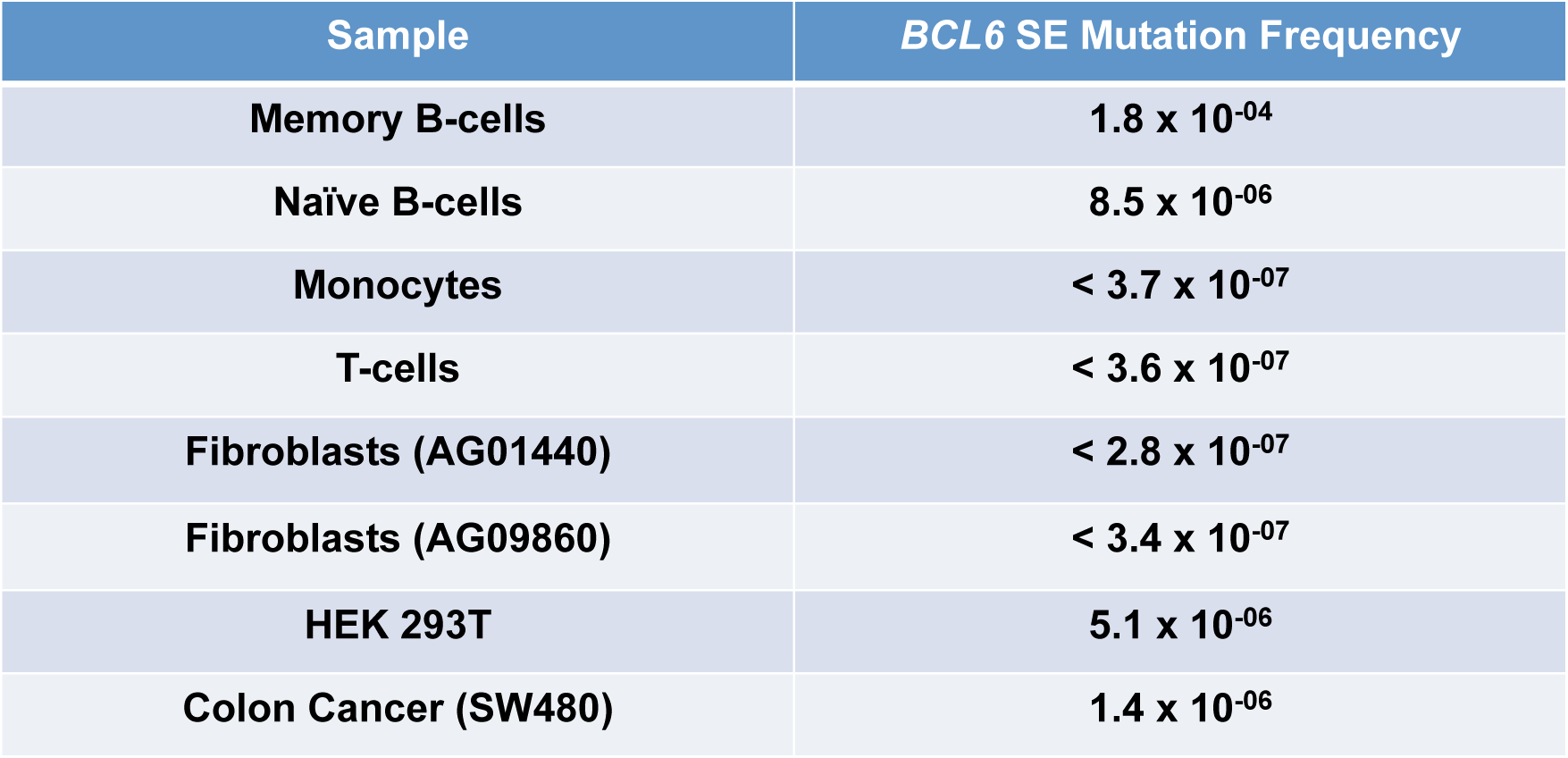
Mutation frequencies of *BCL6* SE in various cell types.

| Sample | *BCL6* SE Mutation Frequency |
| --- | --- |
| Memory B-cells | $1.8 \times 10^{-04}$ |
| Naïve B-cells | $8.5 \times 10^{-06}$ |
| Monocytes | $< 3.7 \times 10^{-07}$ |
| T-cells | $< 3.6 \times 10^{-07}$ |
| Fibroblasts (AG01440) | $< 2.8 \times 10^{-07}$ |
| Fibroblasts (AG09860) | $< 3.4 \times 10^{-07}$ |
| HEK 293T | $5.1 \times 10^{-06}$ |
| Colon Cancer (SW480) | $1.4 \times 10^{-06}$ |

The SEs of *PAX5, POU2AF* and *CD83* also showed high mutation frequencies; however, they are over an order of magnitude lower than that of the *BCL6* SE. The mutation frequencies are 9.7 x 10^-6^, 9.4 x 10^-6^ and 6.9 x 10^-6^, respectively (Fig. 1 and Table S3). The SE of *H2AFX* was the least mutated with an average mutation frequency of 2.8 x 10^-7^, close to background detection levels (Fig. 1 and Table S3). The mutation frequencies of the other sequenced SEs ranged between those of *BCL6* and *H2AFX* SEs (Fig. 1 and Table S3). Since large numbers of mutations lend confidence to data interpretation, we focused our studies on the mutational analyses of the *BCL6* SE locus.

### Mutations in the *BCL6* SE locus show a remarkably similar pattern in all individuals

High-resolution duplex DNA sequencing enabled us to delineate the landscape of subclonal mutations in the *BCL6* SE locus (Figs. 2a and 2b). We found that close to 90% of the ∼8000 mutations are located in a 600 bp region that corresponds to a tentative ‘open’ chromatin region enriched for histone markers H3K4me3 and H3K27Ac in germinal center B-cells (20, 21). The density of mutations is extensive with as many as 3 mutations present in a single read (Fig. S1). Within this 600 bp region, we observed two major mutational clusters, each spanning ∼50 nt (Fig. 2a and Fig. S2). The average mutation frequencies for clusters 1 and 2 are 7.2 x 10^-4^ and 7.7 x 10^-4^, respectively. These contrast a frequency of 3.5 x 10^-6^ in a 50-bp region downstream from the hotspot clusters (Fig. 2a). The mutation load at the two hotspots suggests that 20-80% of memory B-cells possess at least one mutation within one of these clusters. Notably, both the location of these mutation clusters as well as the mutation frequencies are remarkably conserved in healthy individuals across different ethnicities (Figs. 2b and S3; Table S4). The average mutation frequency of the *BCL6* SE is 1.5 x 10^-4^, 2.0 x 10^-4^, 4.2 x 10^-4^ and 2.0 x 10^-4^ in Caucasian, African American, Asian, and Hispanic individuals, respectively (Fig. S3 and Table S4), and the average across all individuals is 2.2 x 10^-4^ (Fig.1 and Table S3). There is no significant difference among the four ethnic groups (*p*=0.26, Fig. S3). We also identified a third cluster of mutations in the TAA/WA sequences, with an average mutation frequency of 5.4 x 10^-4^ (Fig. 2b). Although mutations here are present at high density, they are at lower frequencies than those in clusters 1 and 2. Clustered mutations were also detected in the *PAX5* and *CD83* SEs (Figs. S4a and S4b, respectively), suggesting that SEs of B-cell lineage-specific genes are targeted by a similar mutagenic mechanism.

To confirm that these clustered mutations are specific to the memory B-cell population, we purified and sequenced different cells – memory B-cells, naïve B-cells, monocytes, and T-lymphocytes, as well as cultured primary fibroblasts, transformed embryonic kidney cells (293T) and the colon cancer cell line SW480, using the same SE capture probe set. We found that amongst all cell types, only memory B-lymphocytes have an increased frequency of mutations at the *BCL6* SE locus (Table 1).

### The *BCL6* SE locus is targeted by AID and error-prone DNA polymerases

To gain insights into mutational processes operating at the *BCL6* SE, we examined the spectrum of base substitutions. We observed that transition base-substitutions occur more frequently than transversions (Fig. S5a). A>G/T>C and C>T/G>A transitions account for 28% and 23%, respectively, of all mutations, while C>G/G>C, A>C/T>G, A>T/T>A and C>A/G>T transversions constitute 18%, 13%, 12%, and 4%, respectively, of the remainder. The frequency of transition base-substitutions in the *BCL6* SE is similar to that observed at the SHM target locus *IgHV3-*23, in which C>T/G>A and A>G/T>C transitions comprise 34% and 23%, respectively, of all mutations (Fig. S5b). The C>T transitions in SHM are primarily generated by AID. AID is an initiator of SHM; it deaminates cytidine to uracil, which base pairs with adenine. In addition to AID, error-prone DNA polymerases are also implicated in SHM. The leading candidate is Pol η. Of the known error-prone DNA polymerases, Pol η is believed to function in a non-canonical base excision repair (BER) or mismatch repair (MMR) pathway following AID-catalyzed cytidine deamination (4). Pol η preferentially generates G:T or A:C mismatches at A:T base pairs during DNA synthesis to generate A>G/T>C transition mutations (22). This is what we observed – almost every A/T site within the TAA/WA repeat region in the *BCL6* SE shows a predominance of transition mutations (Fig. S6). Our mutation spectrum data suggest that, similar to the immunoglobulin genes, the *BCL6* SE locus is a target of AID and Pol η resulting in a high prevalence (>60%) of A>G and C>T transition mutations. Moreover, the C>G/G>C transversions we observed could also result from the sequential activities of AID and error-prone DNA replication. The G:U mismatches generated by AID can be efficiently converted to G:apurinic/apyrimidinic (AP) nucleotide pairs by uracil DNA glycosylase (UDG). While replicative DNA polymerases insert A residues across unrepaired AP sites (23), error-prone polymerases such as Rev1 preferentially insert C residues to generate C>G/G>C transversions instead of C>T/G>A transitions (24, 25). Pol η may also play a role in AID-induced C>G transversions (24, 25) but the mechanism is still not clear. Thus, the contribution of AID, Rev1 and Pol η to mutations (C>T, C>G and A>G) within the *BCL6* SE could be as high as ∼70% (Fig. S5a).

### Mutant allele fractions reveal the order of mutagenesis at the *BCL6* SE

Allele fractions of mutations can be used to unravel the order of mutagenic events in a cell population. In this case, higher allele fractions would reflect earlier events and lower fractions, more recent events. Assuming this is true for mutations at the *BCL6* SE locus, we delineated the order of mutagenic events by parsing the observed mutations into 5 bins corresponding to the following mutant allele fractions (MAF) – bin 1: >1x10^-3^, bin 2: >5x10^-4^-1x10^-3^, bin 3: >1x10^-4^-5x10^-4^, bin 4: >5x10^-5^-1x10^-4^ and bin 5: >1.7x10^-5^-5x10^-5^. We then determined if there were differences in the spectrum of base substitutions in different bins.

The base changes at C residues, resulting primarily from the activity of AID are likely the earliest events as C>T/G>A and C>G/G>C alterations comprise >70% of all mutation types in bin1 (highest MAF; Figs. 3a and S7a). A>G/T>C substitutions, reflective of Pol η activity, increase in abundance in bins 2 and 3 concomitant with a decrease in C substitutions. A>G/T>C substitutions then decrease in bins 4 and 5 while C>T/G>A and C>G/G>C alterations increase progressively. A similar pattern was observed when we compared substitutions at G:C *versus* A:T pairs. Close to 80% of the substitutions in bin 1 are at G:C pairs; this fraction decreased in bins 2 and 3 but then increased in bins 4 and 5 in coordination with an increase and decrease, respectively, of mutations at A:T pairs (Figs. 3b and S7b). These data suggest that mutations at the *BCL6* SE locus are not the result of random events, but instead arise from the coordinated action of AID and Pol η. They further suggest that AID inscribes its mark on the mutational landscape prior to Pol η. By contrast, mutational footprints of both AID and Pol η appear early (bin 1: MAF >1x10^-3^ – indicative of clonal expansion) in the immunoglobulin gene *IgHV3-23* (Fig. S8). After initial rounds of selection, however, AID catalysis becomes predominant in bin 2 (MAF >5x10^-4^-1x10^-3^), with approximately equal contributions of AID and Pol η thereafter (Fig. S8).

### Mutational signatures of *BCL6* SE are consistent with the contribution of AID, Pol η **and Rev1 to aSHM**

We determined mutational signatures at the *BCL6* SE by examining the identity of adjacent 5’- and 3’- nucleotides (26). We observed that A>G/T>C transitions are mostly present at A<u>T</u>A sites (Fig. 4a), while both C>T/G>A transitions and C>G/G>C transversions most often occur in the tri-nucleotide sequence, A<u>C</u>A (Fig. 4a). This spectrum is similar in all ten B-cell samples (Fig. S9a). In contrast, C>T/G>A transition mutations in *IgHV3-23* show a sequence preference for G<u>C</u>T and A<u>C</u>C sites, whereas C>G/G>C transversions are more frequent at G<u>C</u>T and A<u>C</u>T sequences (Fig. 4b). Moreover, mutational signatures of *IgHV3-23* differ among the 10 individuals (Fig. S9b), likely due to selection of different mutant *IgH* clones in each person.

We further used deconstructSigs (27) to deconstruct our signatures into COSMICv3 Single Base Substitution (SBS) signatures. We identified six mutational signatures in *IgHV3-23* (Fig. 4b). Two of these signatures-SBS 84 (25.2%) and SBS 85 (9%) have SHM-related etiology. SBS 84 shows predominant C>T transitions (likely AID-mediated) and SBS 85 has equal contributions of T>A and T>C substitutions (likely the result of DNA synthesis by error-prone DNA polymerases). We extracted two additional signatures, SBS 37 and SBS 39, which also could potentially be attributed to error-prone DNA polymerase activity during SHM. SBS 37 (9%) and SBS 39 (23.3%) are represented by T>C transitions and by C>G transversions, respectively (Fig. S10). Similarly, we identified six mutational signatures at the *BCL6* SE (Fig. 4a); three of these, SBS 85 (8.6%), SBS 37 (13.1%), and SBS 39 (28.6%), are shared with *IgHV3-23*. While the etiology of SBS85 is known, those of SBS 37 and SBS 39 are not. Based on the spectrum of nucleotide substitutions found in SBS 37 and SBS 39, and their presence in *IgHV3-23* (the target of SHM), we hypothesize that SBS 37 and SBS 39 could be signatures of error-prone DNA polymerases in SHM. A combination of signatures SBS 37, 39, 84, and/or 85 was also identified at the *PAX5* and *CD83* SE loci where they accounted for as much as 75% of all signatures (Fig. S11). Thus, signatures of SHM permeate the mutational landscape of B-cell SEs.

## Discussion

We sequenced the SE regions of 12 highly expressed genes in B-cells from 10 healthy individuals. Previous studies of mutagenesis of B-cell SEs were based on low-resolution single-cell PCR sequencing (8) or mutant mouse models (5). A recent study involving whole genome sequencing of single human B-cells reports mutations at the *BCL6* SE locus. However, very few mutations were identified in this study precluding a detailed analysis of aSHM (28). We utilized exceptionally accurate, high-depth duplex sequencing (15, 16) to reveal a high-resolution landscape of mutations and mutational processes in B-cell SEs in normal healthy individuals.

We show that, of all sequenced SEs, the *BCL6* SE is the most favored target of aSHM with ∼8000 subclonal mutations in total in 10 individuals. The average mutation frequency of 2.2 x 10^-4^ at the *BCL6* SE locus is as high as the SHM frequency of the immunoglobulin genes (Fig. 1 and Table S3). The accurate identification of large numbers of single nucleotide substitutions at the single-nucleotide level enabled the identification of refined clustered mutations. There are three clusters of mutations within the *BCL6* SE, two of which overlap with AID target motifs (RGYW) (29) and a third that lies within the TAA/WA repeat sequences (Figs. 2b, S2 and S6) (30). Additionally, there are multiple mutations, located <100 nt apart, in multiple sequence reads (Fig. S1). While the density of mutations within these clusters is high, the allele frequency of individual mutations is under 0.5%, and most are even lower (Fig. 2b). Remarkably, the location and allele frequency of these clustered mutations is highly conserved in individuals across different ethnicities (Fig. 2b and Table S2). *BCL6* and *IgH* could be within the same topologically associating domain or proximal to each other in the chromatin architecture (7) therefore, the subclonal mutations in the *BCL6* SE may be the result of collateral damage of SHM at the *IgH* locus. The unusually high numbers of mutations at the *BCL6* SE locus suggests that they are not random events, but rather, may be a part of the normal processes of B-cell maturation and differentiation.

The identification of thousands of mutations in the *BCL6* SE enabled the extraction of mutation signatures and analyses of mutational processes operative during aSHM in normal cells. We show that mutations at the *BCL6* SE are dominated by A>G, C>T and C>G substitutions (Fig. S5a). Together, these three mutation types comprise 70% of all mutations at this locus. While C>T mutations are in accord with cytidine deamination by AID (4), A>G and C>G substitutions are consistent with synthesis by error-prone DNA polymerases, Pol η and Rev1, respectively (22, 24, 25, 31–34). It is well established that AID is essential for B-cell development in response to antigen stimulation and for antibody diversification in germinal centers (35, 36). Multiple error-prone DNA polymerases have been implicated in SHM (33, 37–39); of these, Pol η in particular shows increased expression in germinal center B-cells (31, 40, 41) and can be visualized specifically expressed in the dark zone – where SHM takes place (42). The activities of both mutators are also evident in the extracted mutational signatures (Fig. 4a). SBS84 and SBS 85 are designated as signatures of AID-induced somatic mutagenesis in lymphoid cells in COSMICv3. We suggest that SBS 37 and SBS 39 are signatures of error-prone DNA polymerases, Pol η and Rev1, respectively, in SHM, based on their prevalence in the SHM target, *IgHV3-23*, and the dominance of nucleotide substitutions characteristic of Pol η (22) and Rev1 (24). This notion is further supported by the finding of a suggestive Pol η signature similar to SBS 37, with prevalence of A>G transitions at WA motifs, in clustered mutations in human B-cell lymphomas (43). SHM thus accounts for the etiology of at least 50% of the mutational signatures at *BCL6* SE. Approximately one quarter of mutations are of unknown signatures (Fig. 4a), raising the possibility that mutagenic processes besides SHM also operate at the *BCL6* SE locus.

Mutant allele frequencies also facilitated an evaluation of the order of mutational events during aSHM. We show that AID initiates aSHM and Pol η operates later to extend the mutational landscape (Fig. 3). While the order of AID and Pol η mutagenesis during SHM varies slightly from that during aSHM, it is evident that a balance of these two mutagenic activities is required for fine-tuning antibody diversity. This argument is further supported by the finding that GANP (germinal center-associated nuclear protein) plays a critical role in modulating AID and error-prone polymerase activities in germinal center B-cells to facilitate the production of high-affinity antibodies (44).

**Figure 3.**
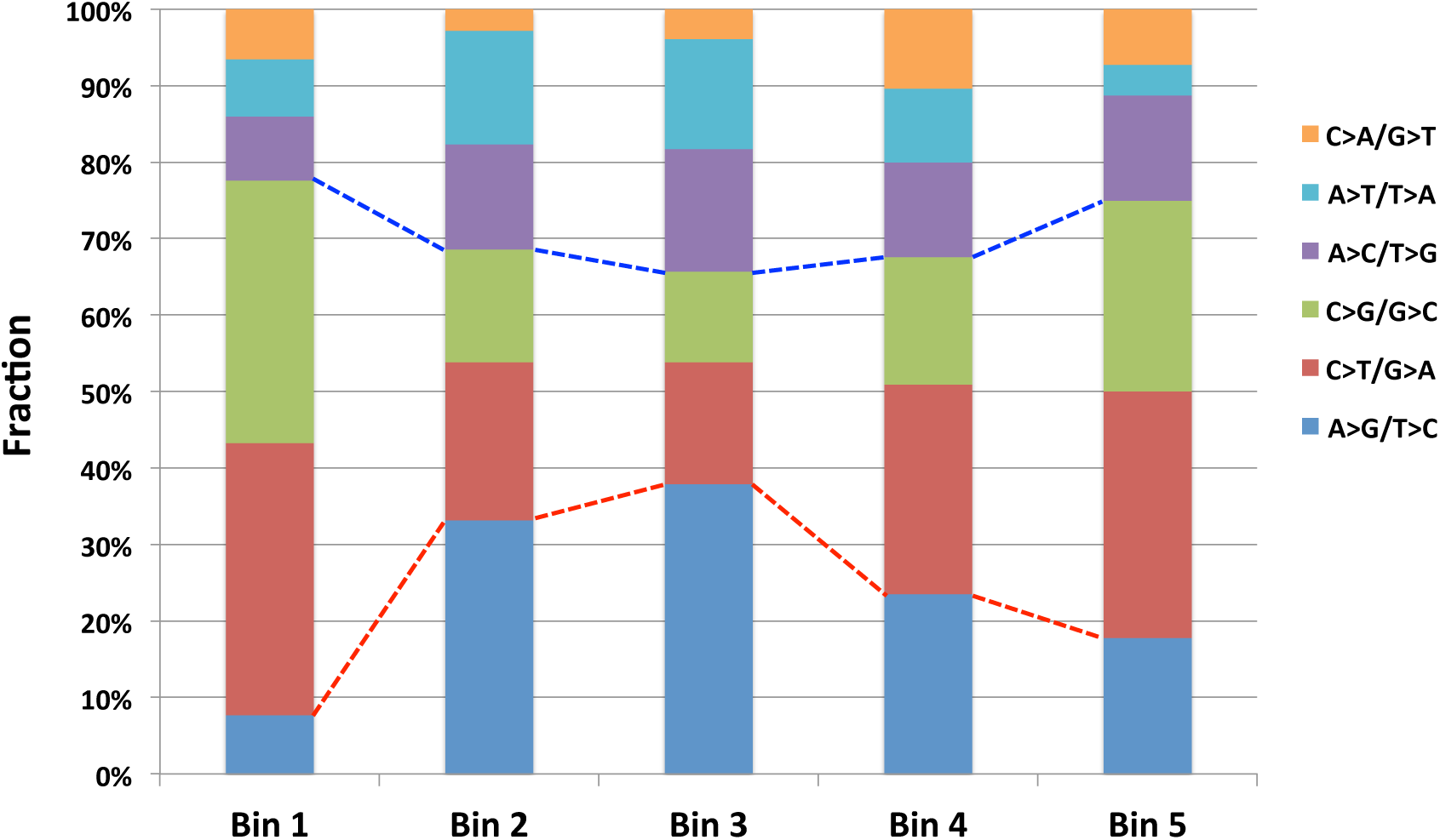
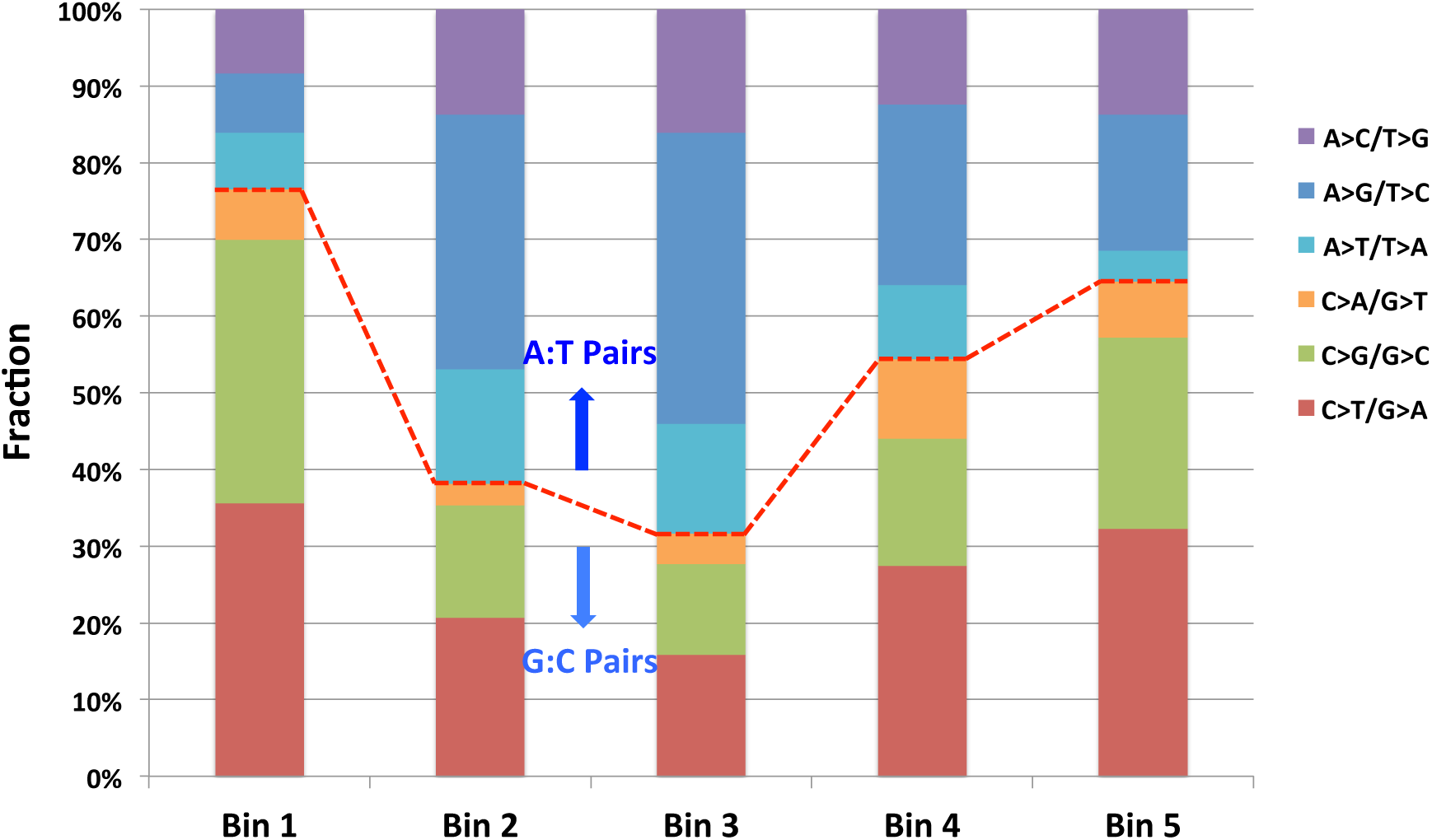
Order of AID and Pol η activities revealed by mutational frequencies and spectra. Cumulative mutations in *BCL6* SE from ten individuals were sorted into five bins based on the mutant allele fractions at each nucleotide position – Bin 1: >1x10^-3^; Bin 2: >5x10^-4^-1x10^-3^; Bin 3: >1x10^-4^-5x10^-4^; Bin 4: >5x10^-5^-1x10^-4^; and Bin 5: >1.7x10^-5^-5x10^-5^. a) Order of mutations as a function of the fraction of each base substitution type in each bin. C>T/G>A (AID-mediated) and C>G/G>C substitutions dominate Bin 1 (∼80%), decrease in Bins 2 and 3, and then gradually increase in Bins 4 and 5. In contrast, A>G/T>C substitutions (Pol η-generated) constitute only a minor fraction of all substitutions in Bin 1, but increase in Bins 2 and 3 before declining. b) Order of mutations at G:C *versus* A:T pairs.

**Figure 4.**
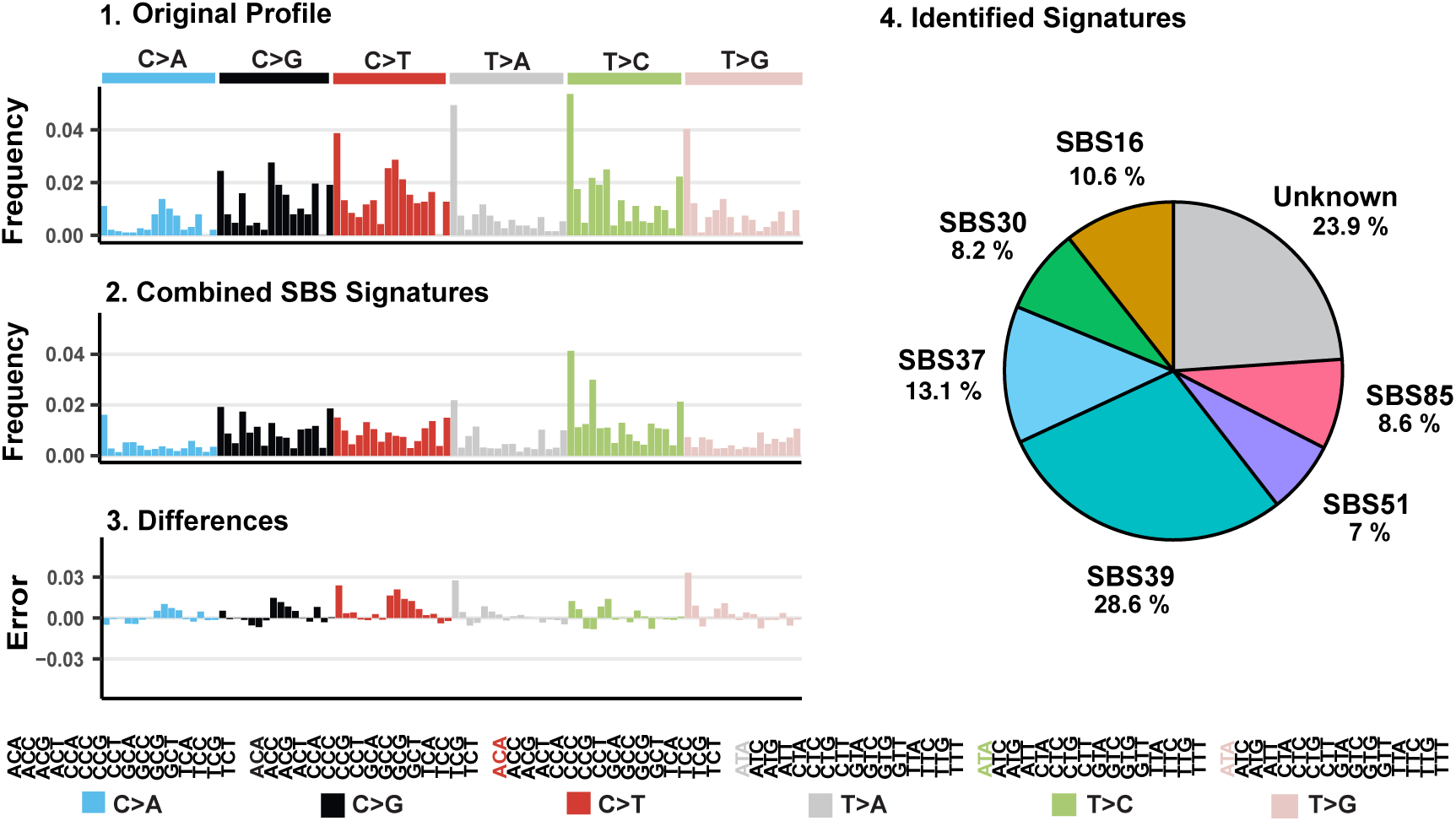
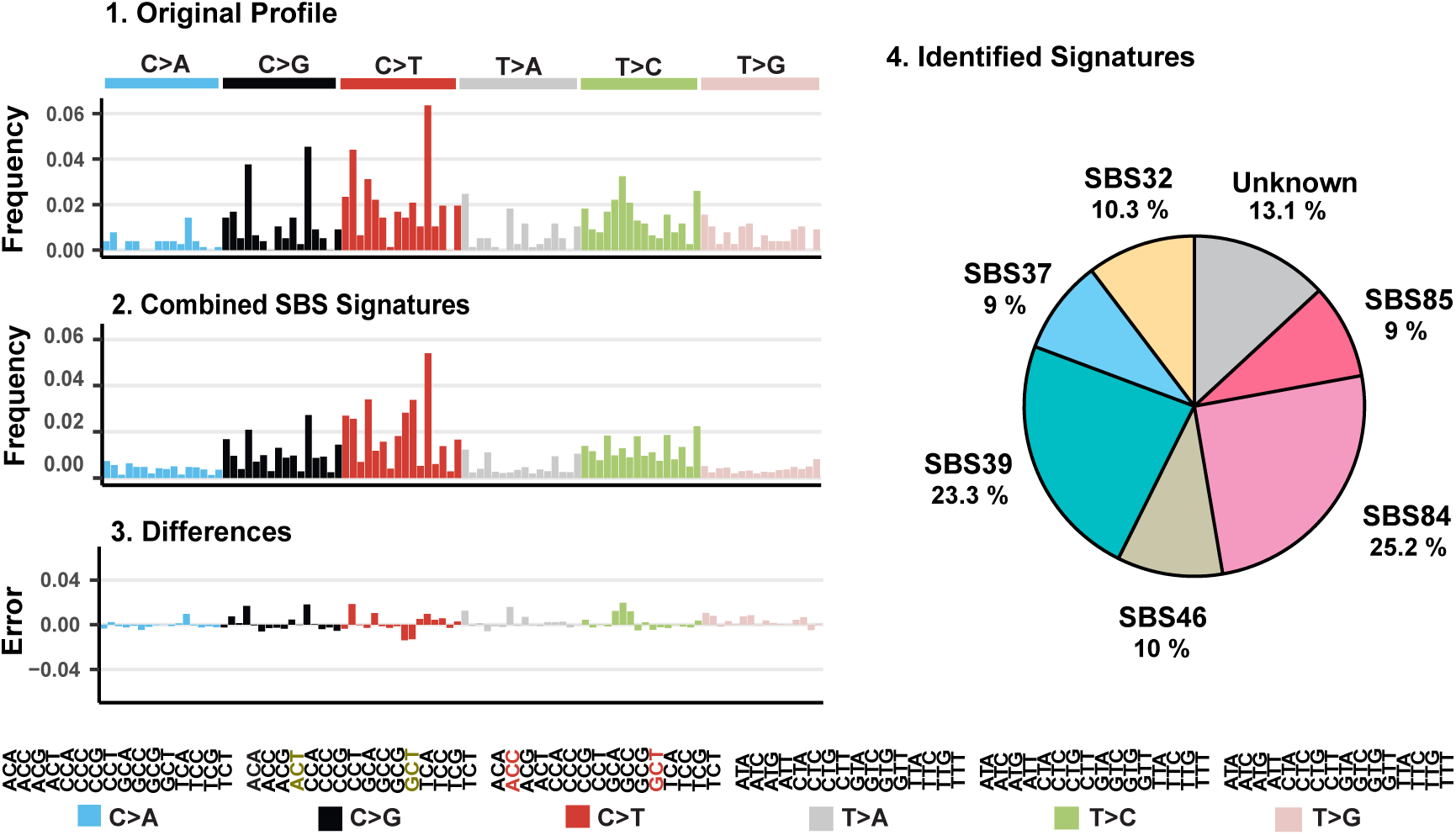
Deconstruction of signatures of unique mutations in *BCL6* SE (a) and *IgHV3-23* (b) from 10 merged B-cell samples using deconstructSigs (24). Panels represent the original profiles (a-1 and b-1), the signature as computed using the best fit combination of COSMICv3 SBS signatures (a-2 and b-2), the difference between the preceding two figures (a-3 and b-3), and the breakdown of the original profile into COSMIC v3 SBS signatures (a-4, b-4). Types of base substitutions and the array of trinucleotide motifs are displayed; motifs with the most frequent substitutions found in the original profile (panel 1) are highlighted with colors designated for each of the six base substitution types.

One of the unique features of the *BCL6* SE mutation landscape is the high density of mutations in DNA sequences located between 400-1,000 bp downstream of the TSS. There is a 20-fold difference in mutation frequencies between sequences with low variations (0-400 bp downstream of TSS; MF 1.7 x 10^-5^) and those with clustered mutations (400-1,000 bp; MF 3.5 x 10^-4^). This demarcation in mutation burden tracks with the positioning of histones H3K27ac and H3K4me3 (21). The “mutation-hot” region shows high levels of histone H3 acetylation and methylation indicative of open chromatin and active transcription with a high density of cis-regulatory elements (45) such as SEs. The cold region is devoid of H3K27ac, suggesting that the nucleosome is in a closed conformation and lacks regulatory function. While an “open” chromatin conformation facilitates active transcription of *BCL6* during B-cell development, it also increases the accessibility of this region to the SHM machinery, which generates mutations and promotes strand breakage (46). Indeed, it has been reported that this “mutation-hot” region is also a hot bed for DNA breaks leading to genomic translocations, frequently fusing *BCL6* with the immunoglobulin heavy chain genes (20, 47).

Multiple rounds of affinity selection and clonal expansion are required for high-affinity antibody production (13). B-cells expressing low affinity immunoglobulins are rapidly eliminated by apoptosis. In the case of *BCL6*, specific subclonal mutations may be selected and expanded to cause aberrant cell differentiation or malignant phenotypes. In fact, clonal *BCL6* SE mutations and translocations are characteristic of diffuse large B-cell lymphomas (12). They can affect the binding of transcription factors such as the IRF4 repressor to deregulate BCL6 protein expression (14); frequently, BCL6 is over-expressed in B-cell lymphomas (48).

The mutational landscape of the *BCL6* SE might serve as a sensitive indicator for mutation loads in healthy individuals. Recent studies have reported the presence of low frequency coding mutations in a large number of genes, including in tumor driver genes, in normal individuals (49–51). Many of these are found clonally expanded in the tumor genome. The B-cell SE subclonal mutations represent a similar class of pre-existent mutations in regulatory regions of genomic DNA. Apparent shifts in the pattern of mutation position, density (the distance between mutations), and/or intensity (the extent of mutations) could signal deregulated gene expression and an enhanced risk for lymphomagenesis.

## Methods

### Human B-cells

The CD19+ B-cells were purchased from AllCells and ReachBio. Peripheral blood mononuclear cells (PBMCs) were purified from whole blood drawn from ten healthy donors (Table S2). CD19+ B-cells were purified by positive selection using an anti-CD19 antibody-conjugated affinity column. Purified memory B-cells (CD27+), naïve B-cells, monocytes, and T-cells, purchased from Bloodworks Northwest, were isolated using cell-specific isolation kits and separation on autoMACS columns (Miltenyl Biotec). All samples were de-identified prior to use.

### DNA isolation

Purified CD19+ B-cells were lysed with proteinase K and DNA was extracted by using the Qiagen DNeasy Blood & Tissue Kit. DNA concentrations were quantified by using the Nanodrop 2000 spectrophotometer.

### Capture library

Synthetic DNA oligonucleotides, purchased from IDT, were employed for targeted gene capture. A capture-probe pool comprising 5’-biotinylated DNA oligonucleotides (120 nt) was designed and used in sequential rounds of hybridization to enrich and sequence the indicated super-enhancers (Table S1).

### Duplex sequencing

Duplex sequencing (DS) was performed using published protocols (15, 16). In brief, genomic DNA from purified CD19+ B-cell was fragmented by sonication, end-repaired and A-tailed. Purified DNA fragments (∼250 bp) were ligated with DS adaptors containing unique molecular barcodes comprising 12 random nucleotides. The ligated DNA was PCR amplified and purified amplicons were hybridized to 5’-biotinylated capture probes. A process of double capture was performed as described previously (52) to improve on-target capture efficiency. After purification by binding to streptavidin beads, the final PCR amplified the captured DNA library using distinct index primers for each sample to enable multiplex sequencing. The prepared libraries were mixed and sequenced by Illumina HiSeq2500.

### Data processing

Sequencing data were processed by previously described methods (16). Sequencing reads with 12-nt molecular barcodes were assembled and aligned to RefSeq to generate single strand consensus sequences (SSCS). Pairs of SSCS with complementary barcodes were grouped to establish double strand consensus sequences (DCS). Mutations were scored only if base substitutions were present at the same position in both strands and were complementary to each other.

## Acknowledgements

We thank Mark Fielden, Herve Lebrec and Hisham Hamadeh for helpful input during the course of these studies, and Clint Valentine for bioinformatics assistance. This study was supported by Amgen, Inc., NIH/NCI Cancer Center Support Grant P30 CA015704, a pilot grant from Core Center of Excellence in Hematology P30 DK 56465, Seattle Translational Tumor Research (STTR) and NIH/NCI P01 CA077852.

## Supplementary Information

### Supplementary Figures

**Figure S1.**
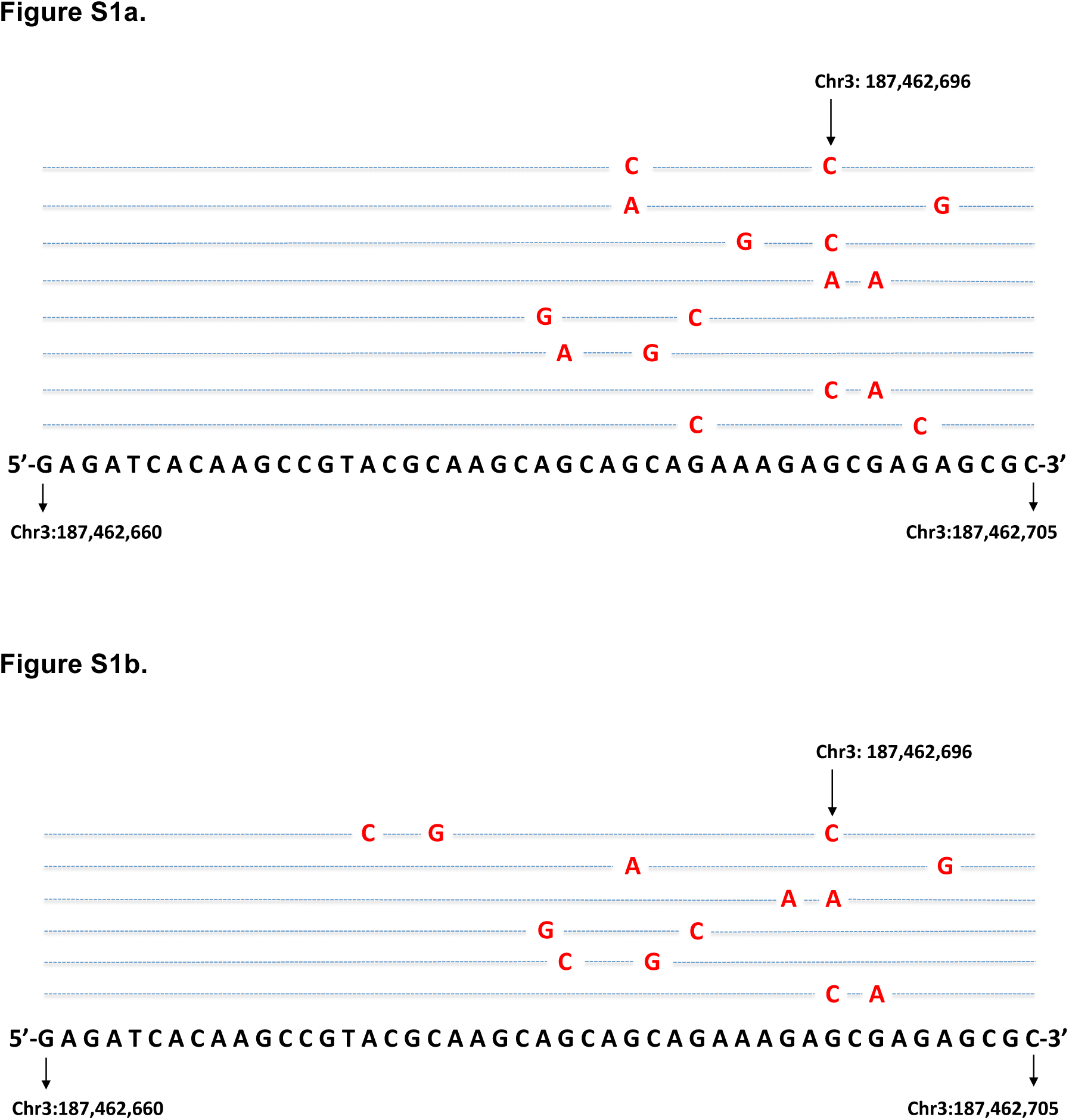
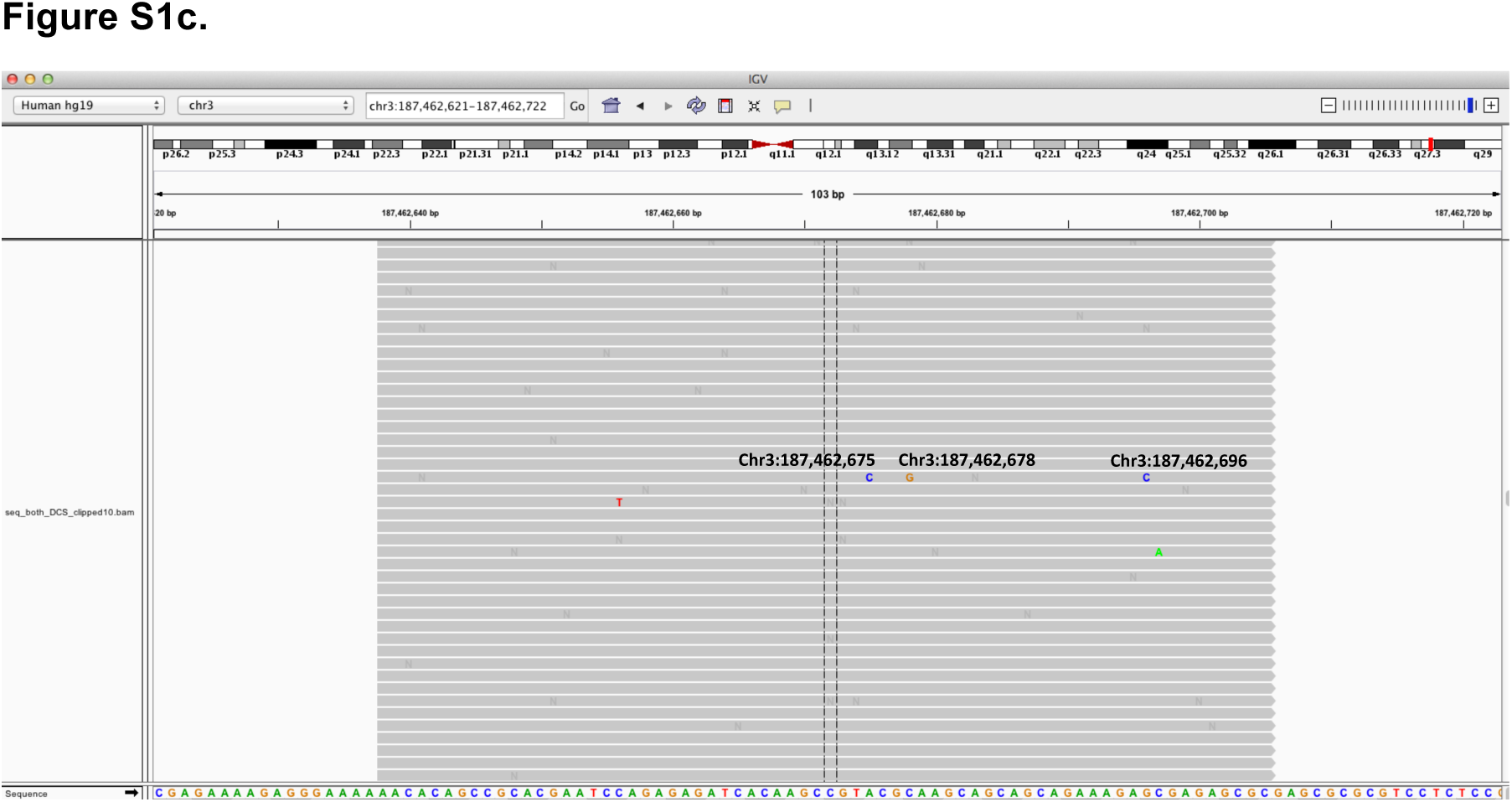
Examples of multiple mutations observed in a single read aligned to the *BCL6* SE locus from one B-cell sample. The duplex sequencing library prepared from sample ID1 was sequenced twice using different indices. Multiple mutations in a single read were observed in both experiments (a) and (b). Up to 3 mutations were observed in a single read in (b). The triple mutations in a single read can be visualized in the IGV graph (c).

**Figure S2.**
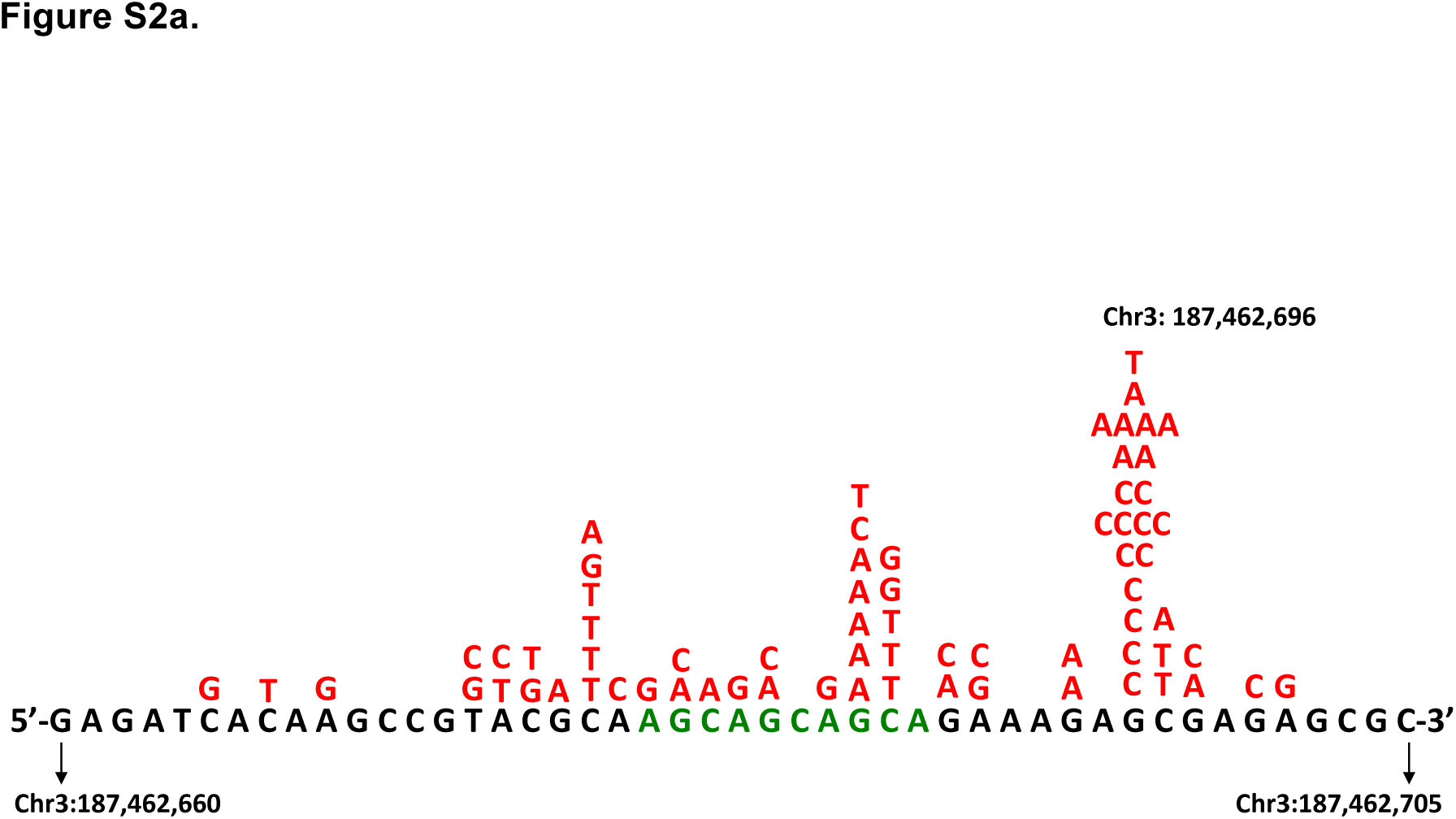
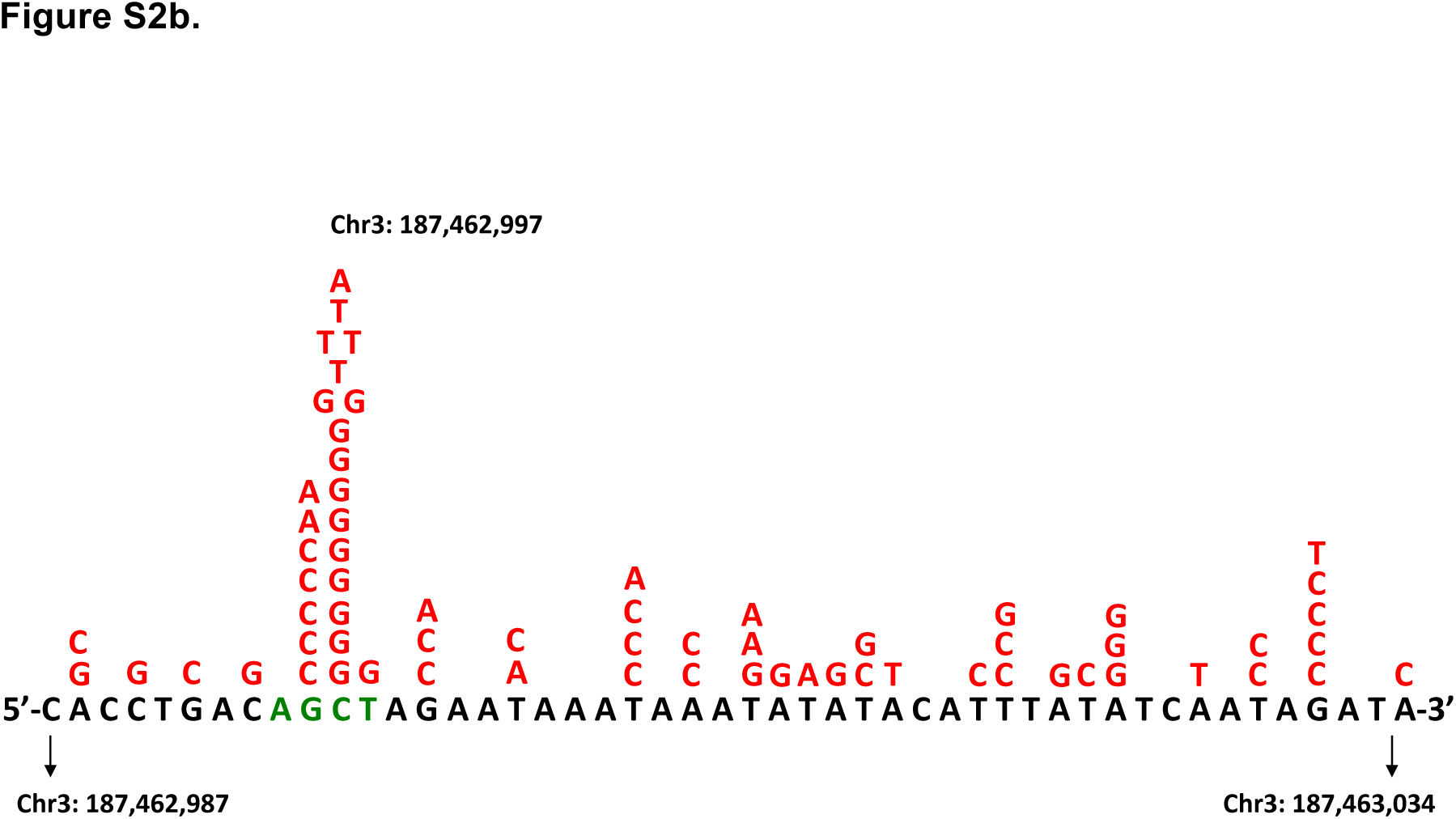
Base-substitution profiles of mutational clusters 1 (a) and 2 (b) in *BCL6* SE in one individual.

**Figure S3.**
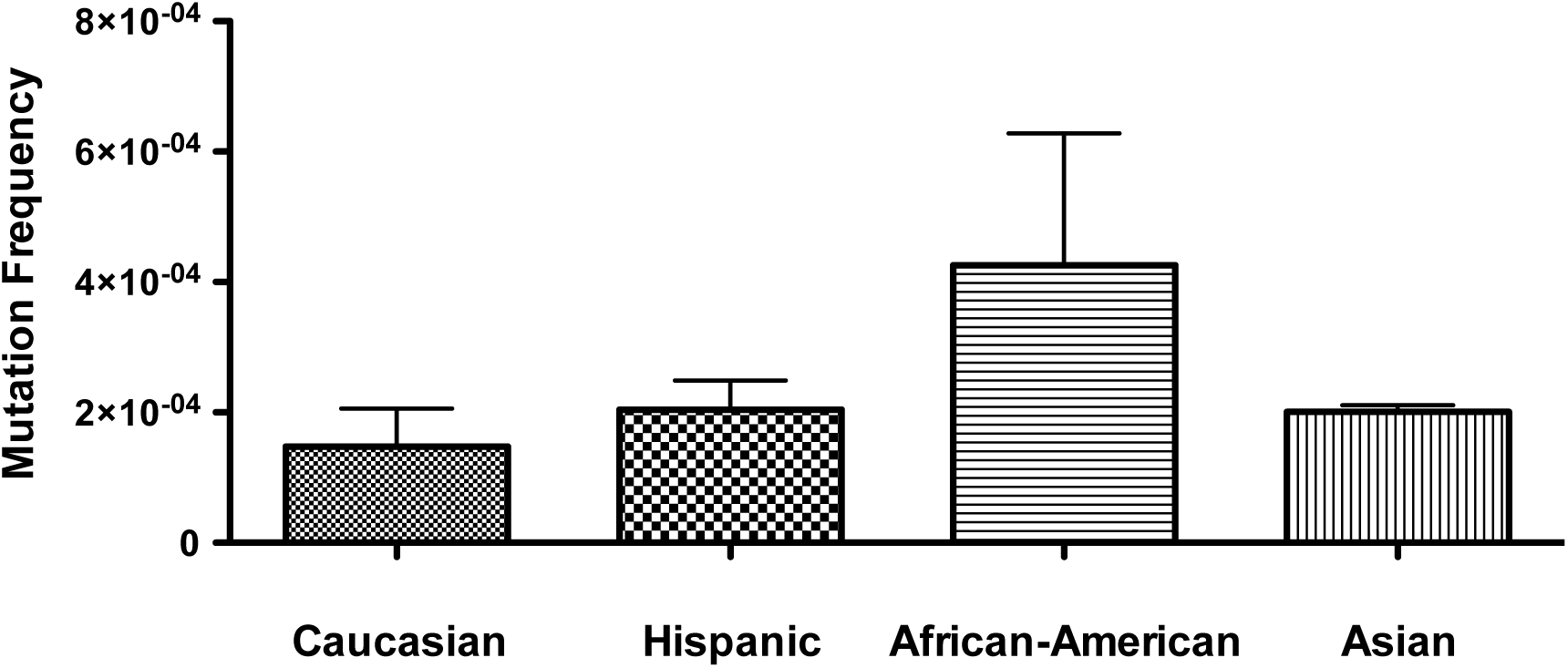
No significant difference in the average mutation frequencies of *BCL6* SE in four different ethnic groups. Mutation frequencies of *BCL6* SE for each individual were calculated and categorized in four ethnic groups: Caucasian, Hispanic, African America and Asian. Data are presented as mean ± SEM (N=3 for Caucasian and Hispanic; N=2 for African American and Asian). There is no significant difference between the four groups as analyzed by ANOVA (*p* = 0.26).

**Figure S4.**
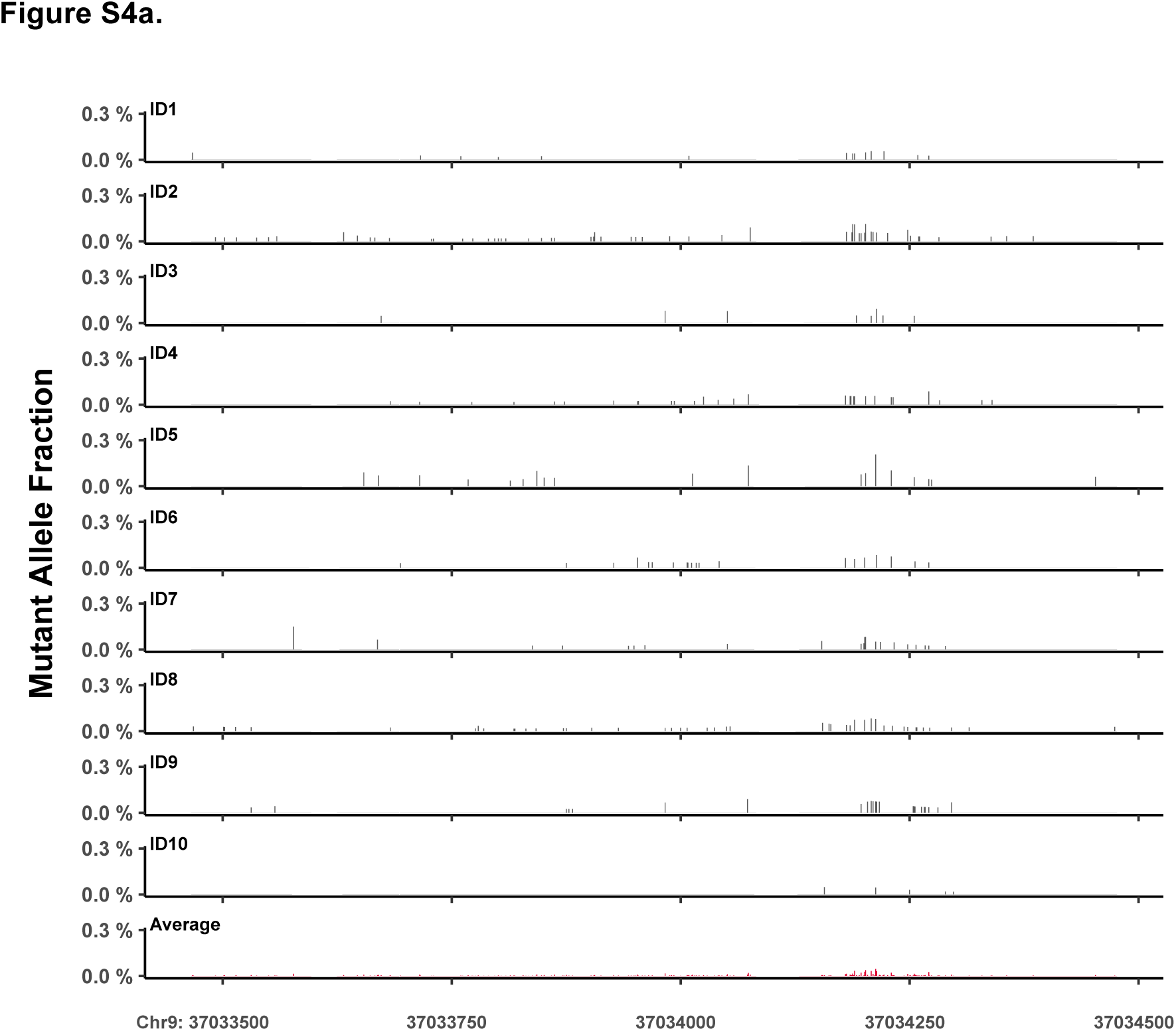
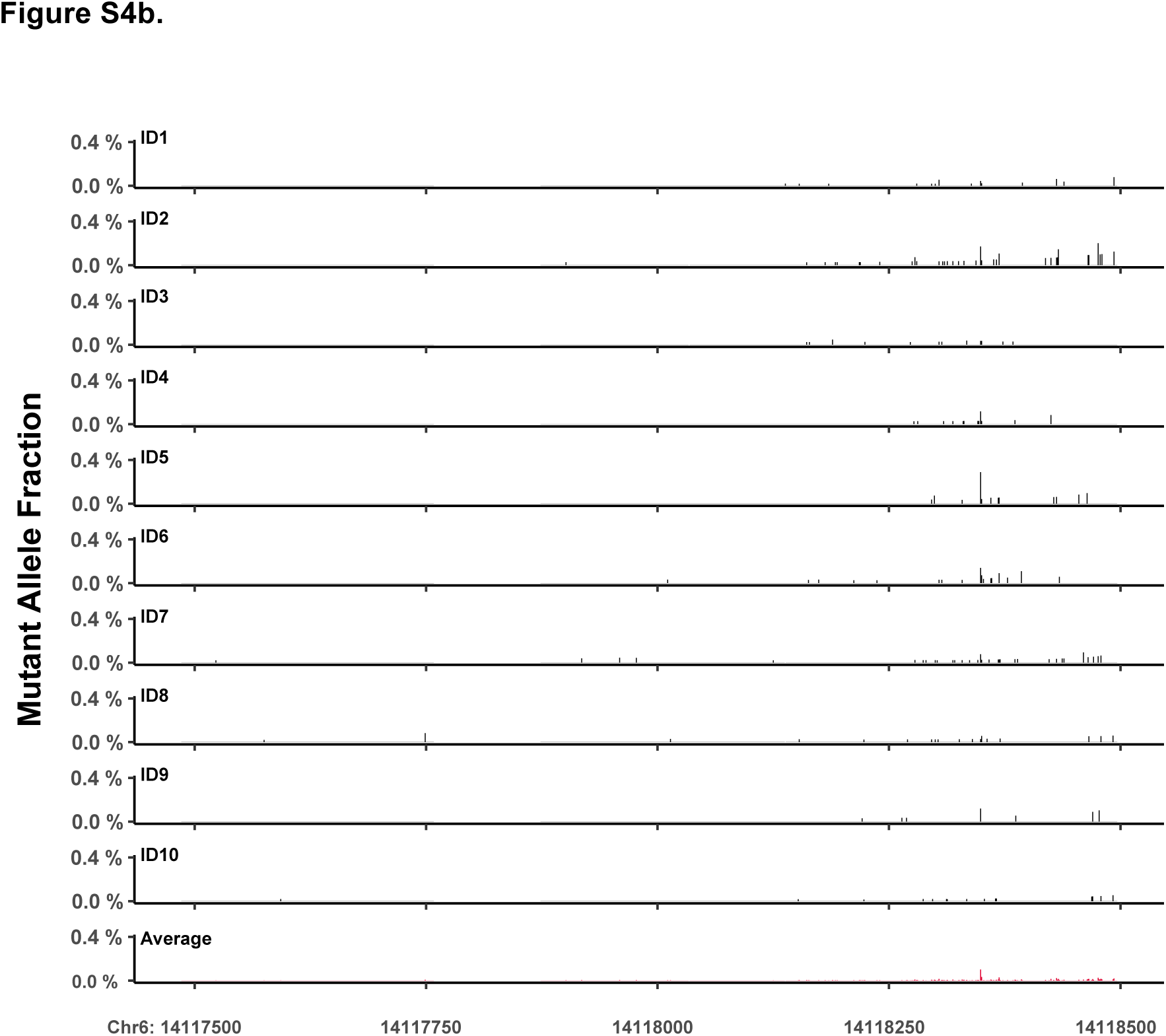
Mutational landscape of *PAX5* (a) and *CD83* (b) SE loci in each of ten individuals. Mutant allele frequencies at each nucleotide position across the sequenced genomic coordinates are presented.

**Figure S5.**
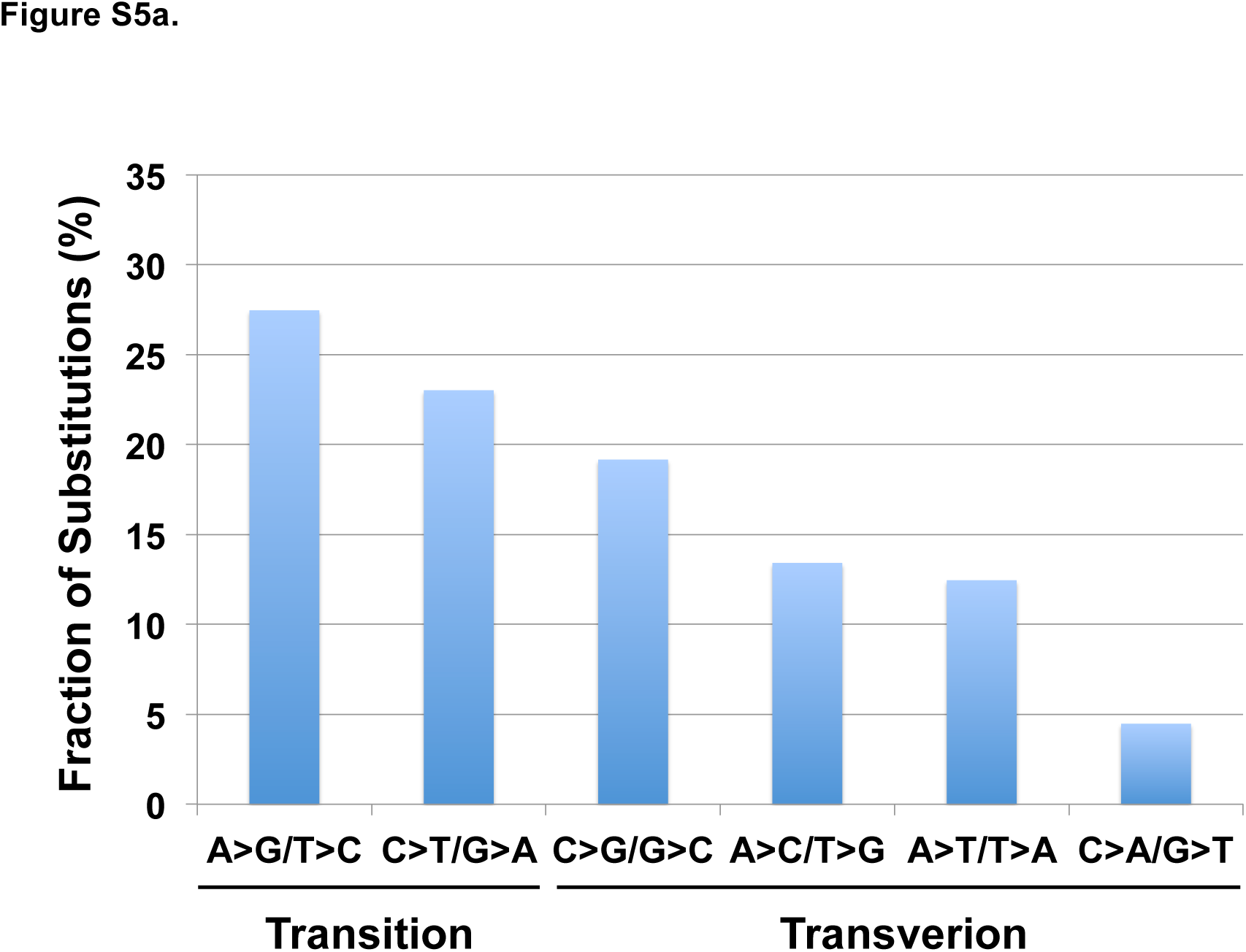
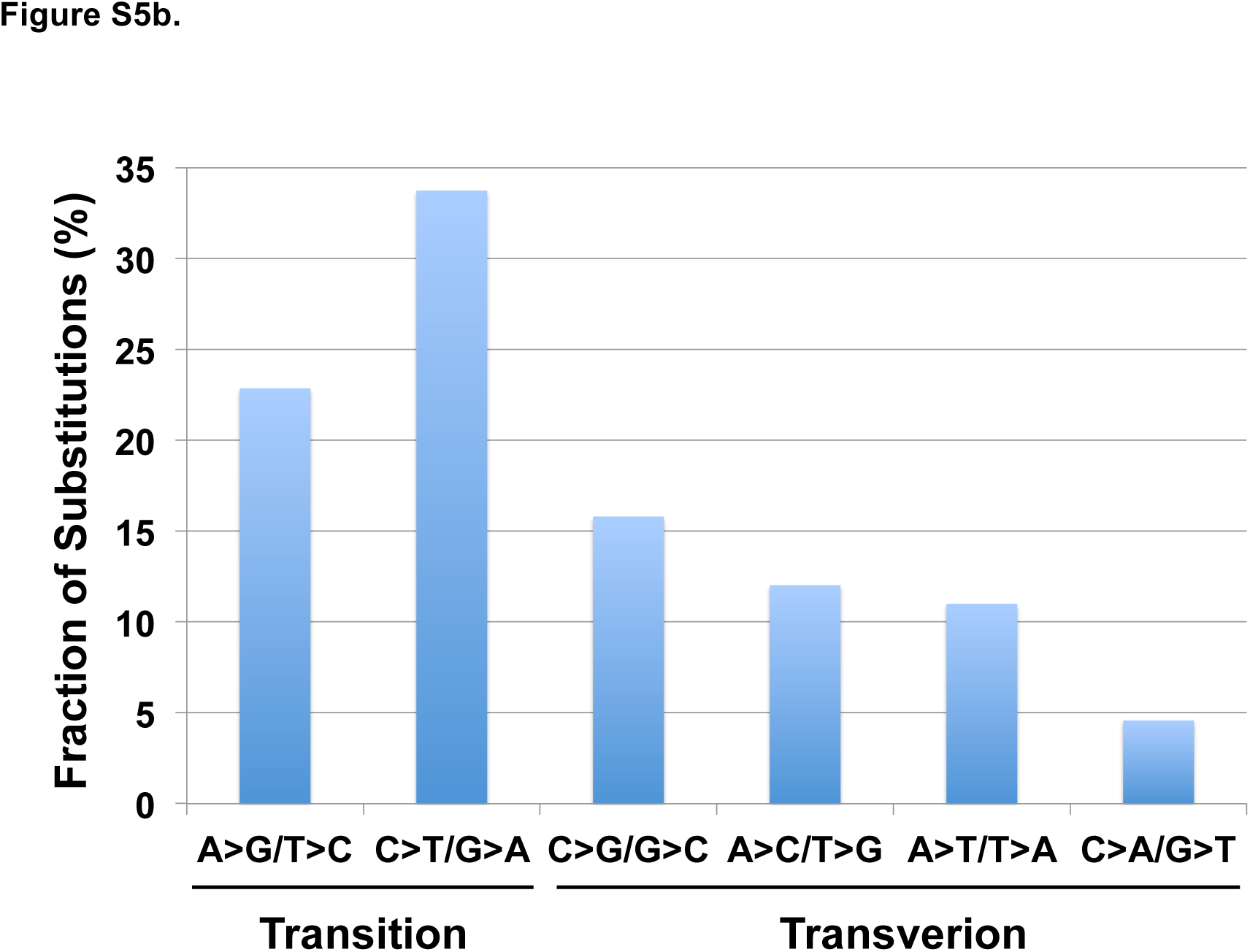
Mutational spectra in *BCL6* SE and in *IgHV3-23*. Fractions of the six possible types of base substitution were determined from a total of ∼8,000 and ∼2,000 mutations detected in the *BCL6* SE and in *IgHV3-23*, respectively. A>G/T>C transitions are most prominent in *BCL6* SE (a) while C>T/G>A transitions are more abundant in *IgHV3-23* (b).

**Figure S6.**
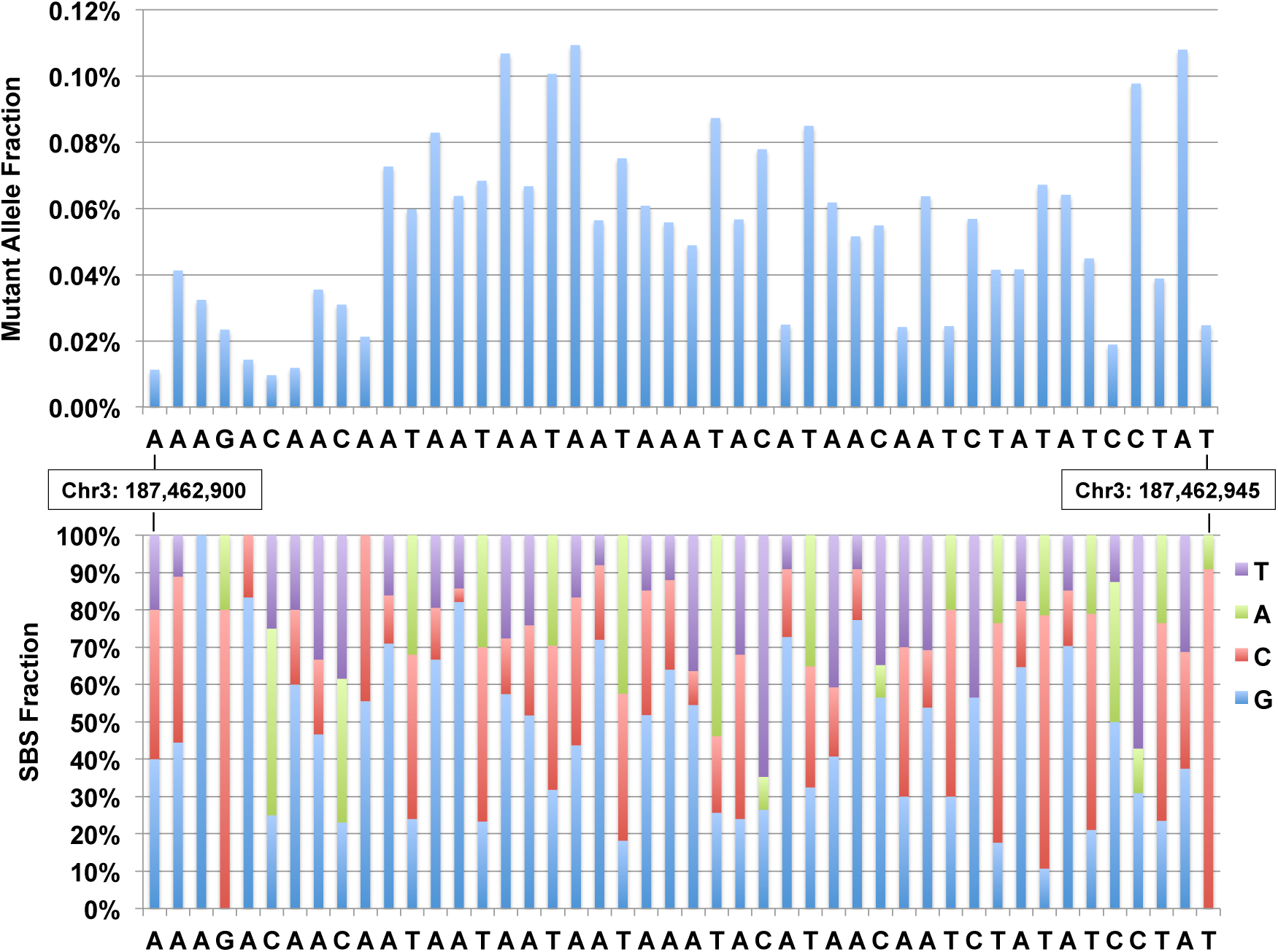
Mutational profile and spectrum in the *BCL6* TAA/WA repeat region. Mutations in this TAA/WA-rich region show dense footprints of the A-T mutator Pol η, as reflected by T>C/A>G substitutions. Although the mutations are clustered at high density, the average mutant allele fraction (∼0.04%) is lower than that in clusters 1 and 2 (∼0.1%, Fig. 2a). No high MAF spikes are observed in this region, contrasting clusters 1 and 2 where the allele fraction of some hotspots is close to 0.5% (Fig. 2a). This discrepancy is likely caused by the fact that only one mutagenic process – error-prone DNA synthesis, functions in the TAA/WA region, while combined AID and error-prone DNA polymerase activities operate at clusters 1 and 2.

**Figure S7.**
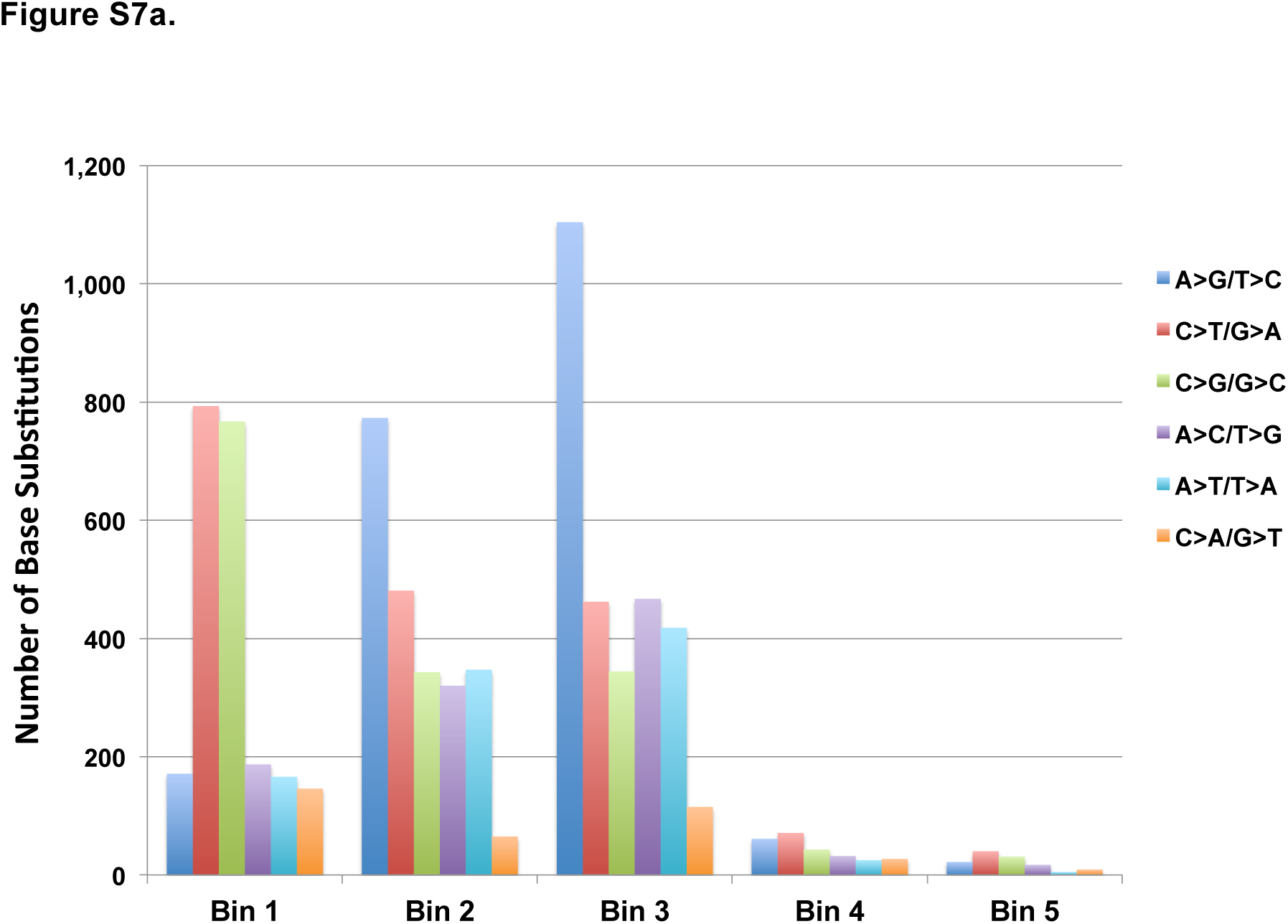
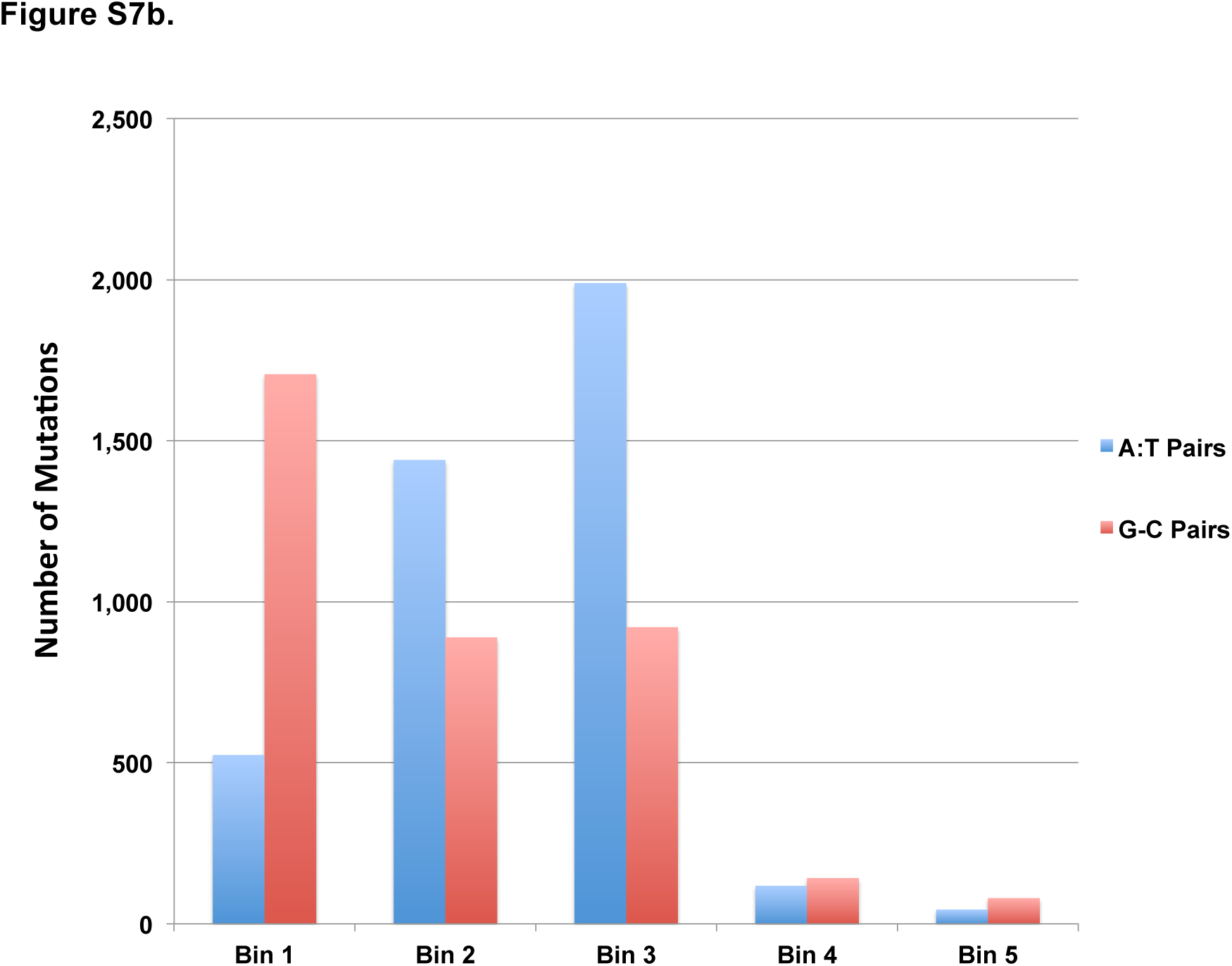
Order of accumulation of base substitutions in *BCL6* SE during aberrant SHM. Mutations in *BCL6* SE from ten individuals were placed into five bins based on the mutant allele fractions – Bin 1: >1x10^-3^; Bin 2: >5x10^-4^-1x10^-3^; Bin 3: >1x10^-4^-5x10^-4^; Bin 4: >5x10^-5^-1x10^-4^; and Bin 5: >1.7x10^-5^-5x10^-5^. a) Numbers of each base substitution type in each bin were calculated and plotted. A>G/T>C transitions increase, whereas C>T transitions and C>G transversions decrease from Bin 1 to Bin 3. b) Numbers of mutations at G:C *versus* A:T pairs in each Bin. The number of mutations at A:T pairs is lower than that at G:C pairs, but gradually increases in Bins 2 and 3. On the other hand, mutations at G:C pair are dominant in Bin 1, but the numbers decrease in Bins 2 and 3.

**Figure S8.**
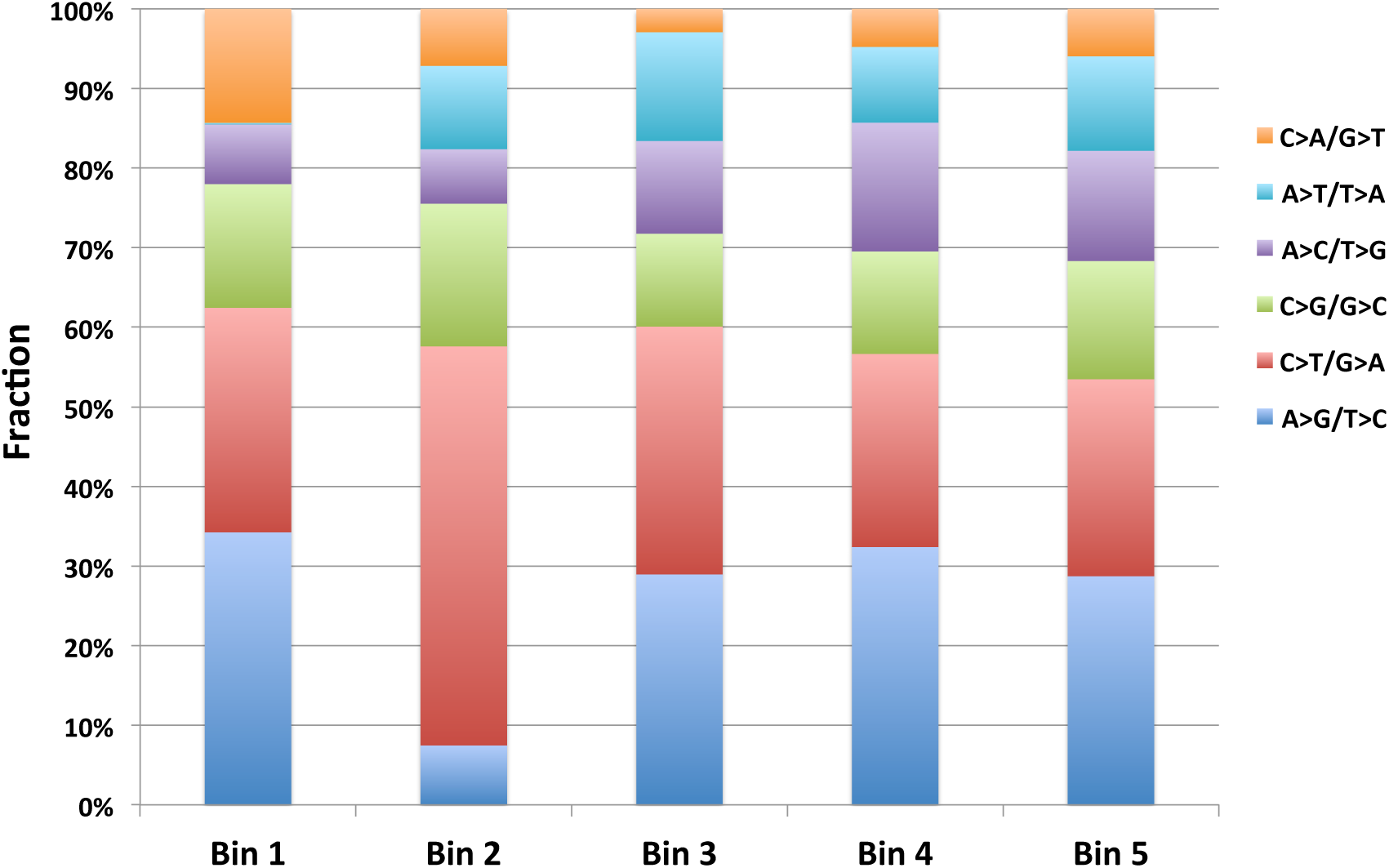
Order of accumulation of base substitutions in *IgHV3-23* during SHM. Order of mutations as a function of the fraction of each base substitution type is plotted as described in Fig. S7 legend.

**Figure S9.**
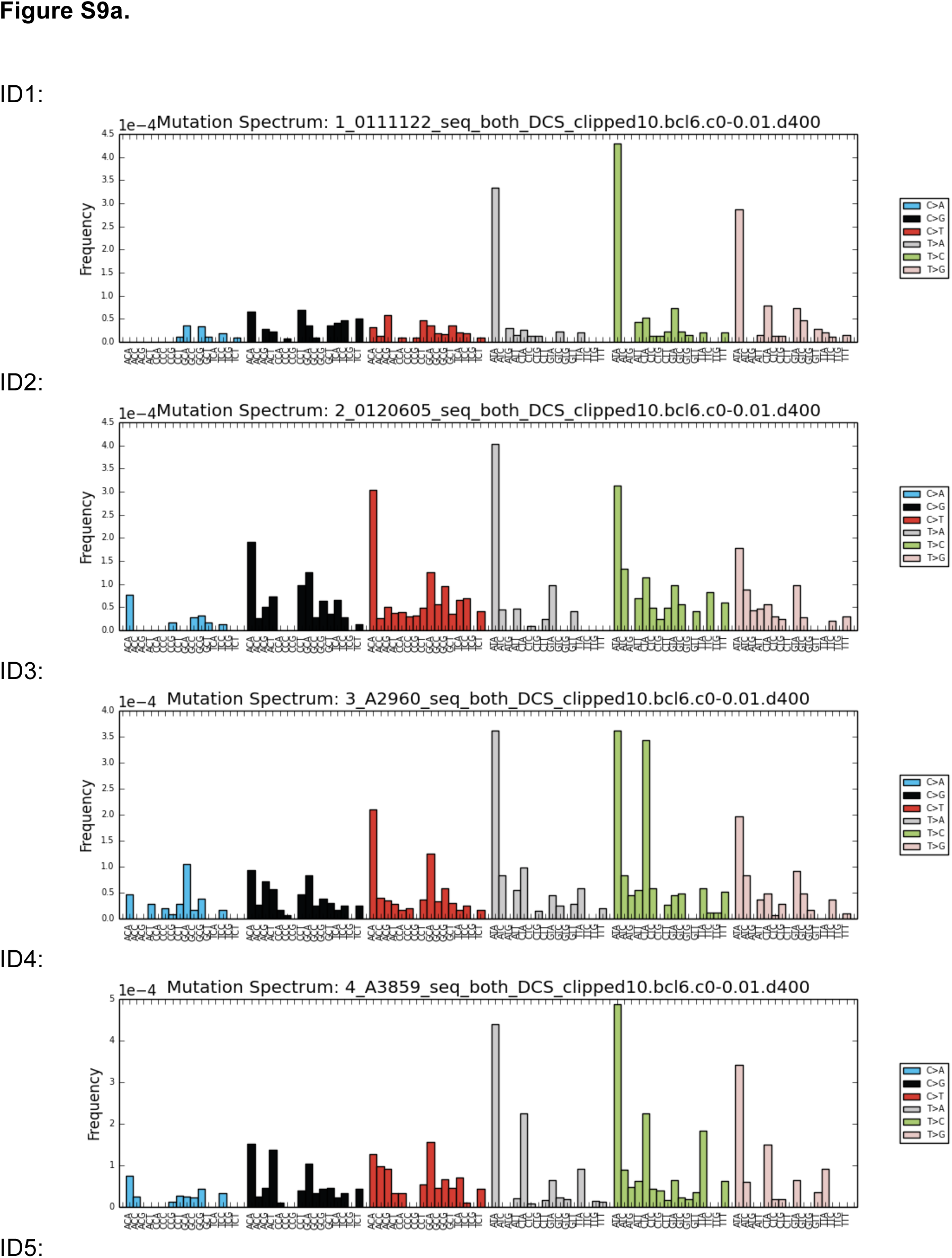
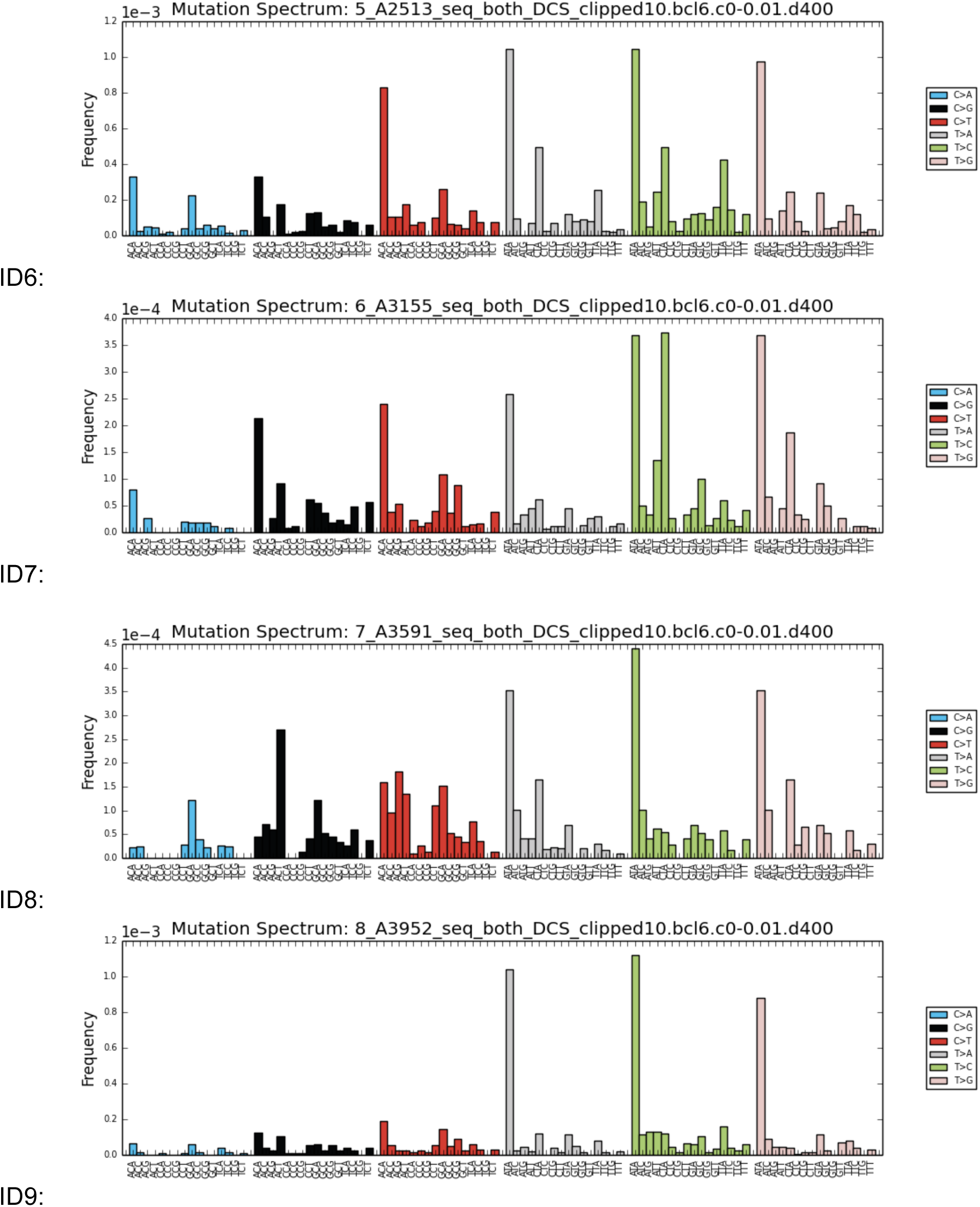
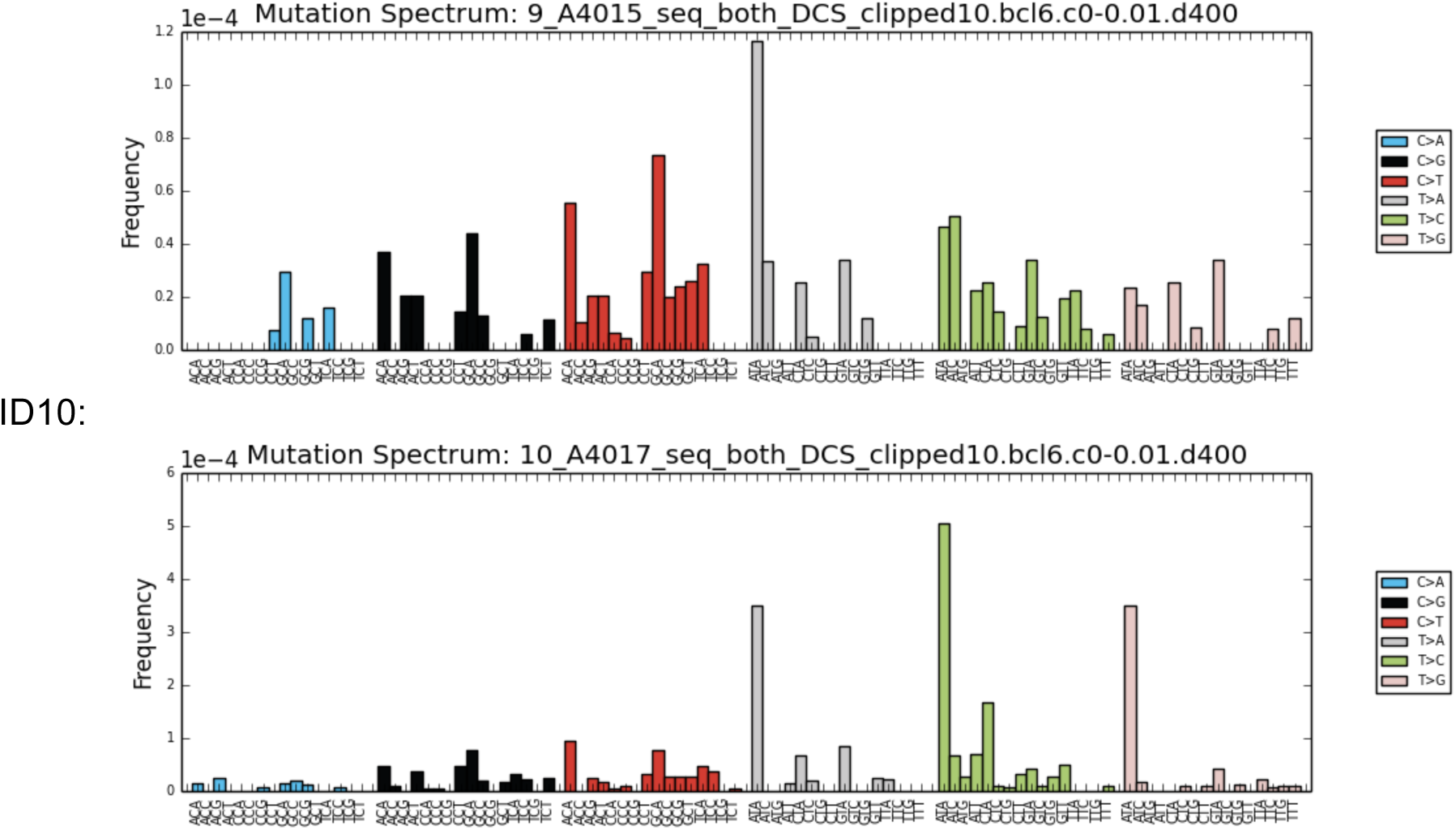
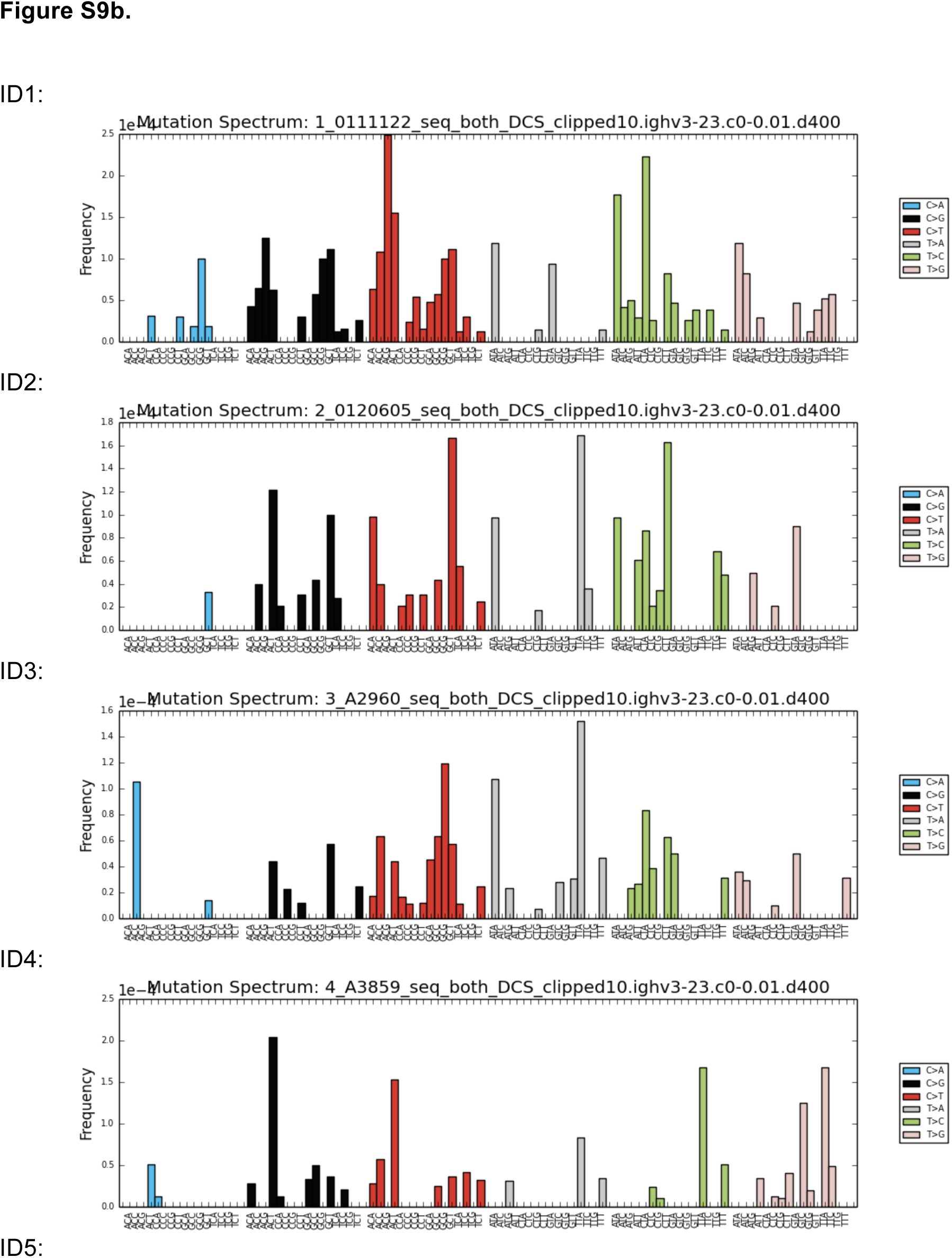
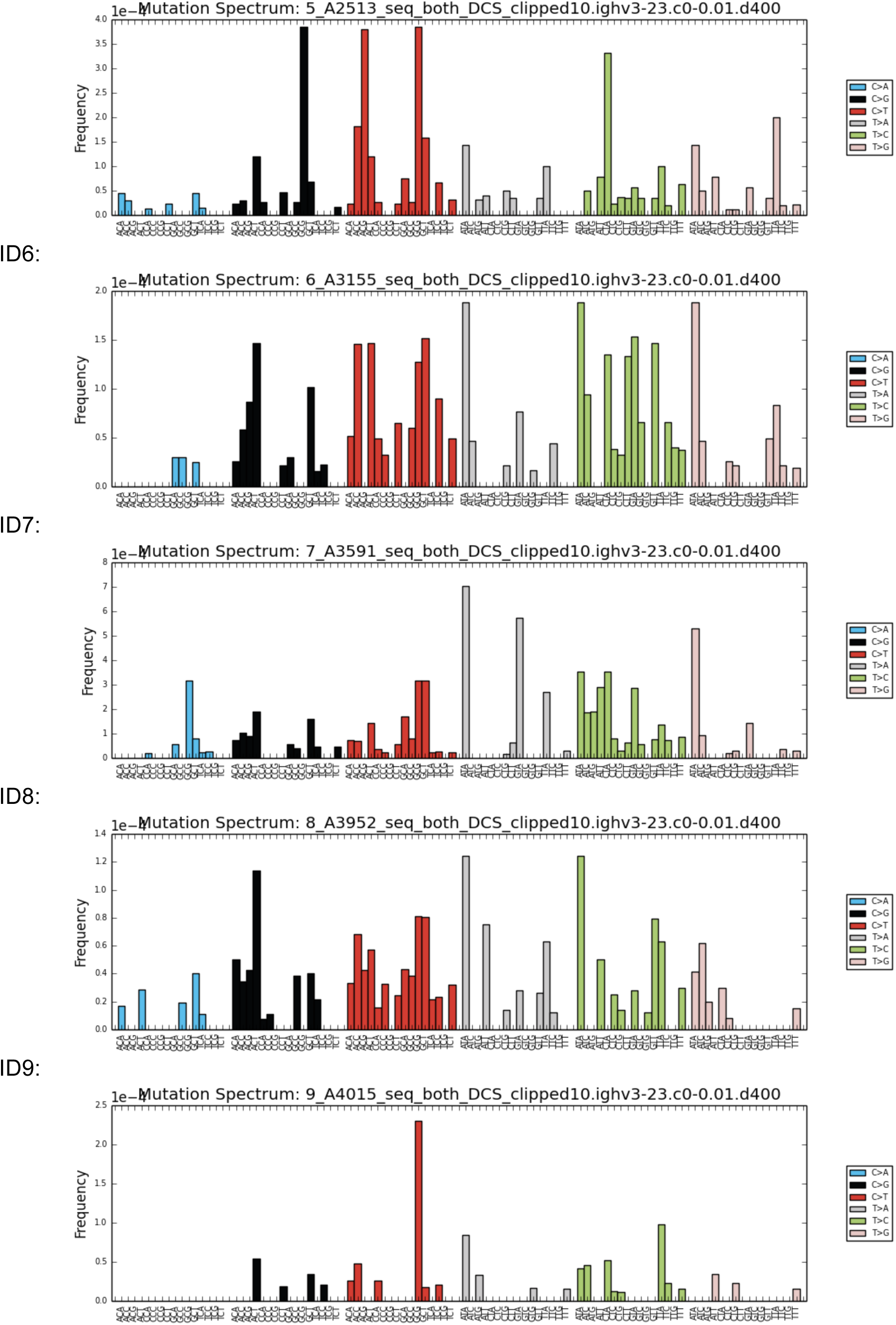
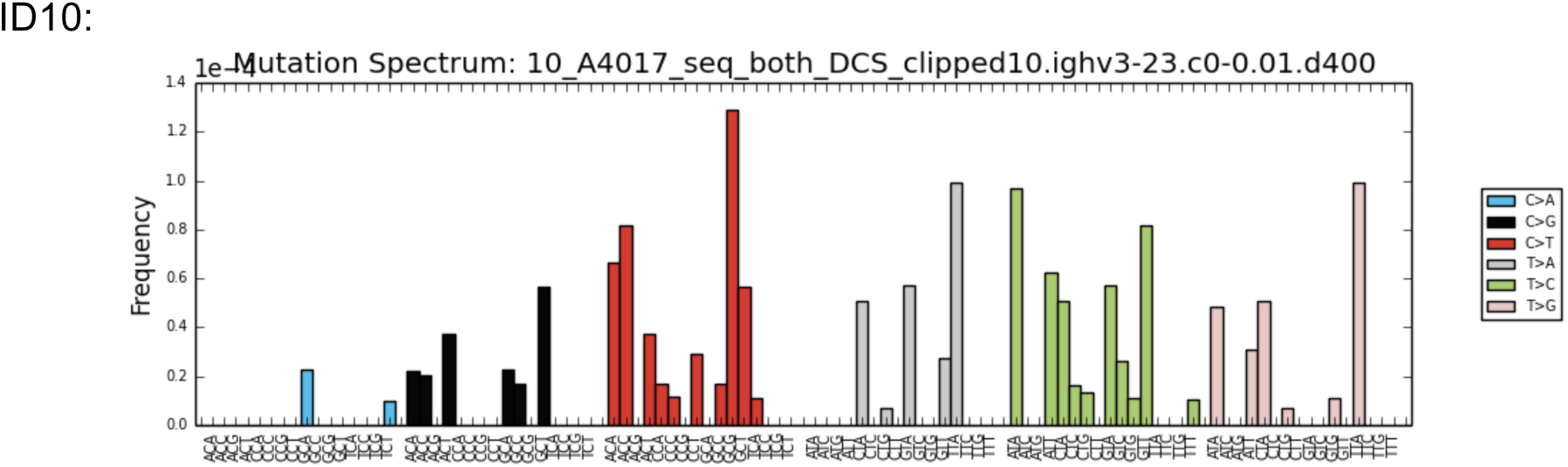
Mutational signatures of *BCL6* (a) and *IgHV3-23* (b) in each individual. The mutational signatures presented in a) indicate profound T>C/A>G substitutions in the tri-nucleotide sequence of A<u>T</u>A, a WA motif preferred by Pol η, and C>T/G>A or C>G/G>C mutations in the G<u>C</u>T sequence – likely in the AID-targeted RGYW motif. These signatures are consistent across different individuals. In contrast, the signatures at *IgHV3-23* (b) differ from each other due to clonal selection of Ig genes in each individual.

**Figure S10.**
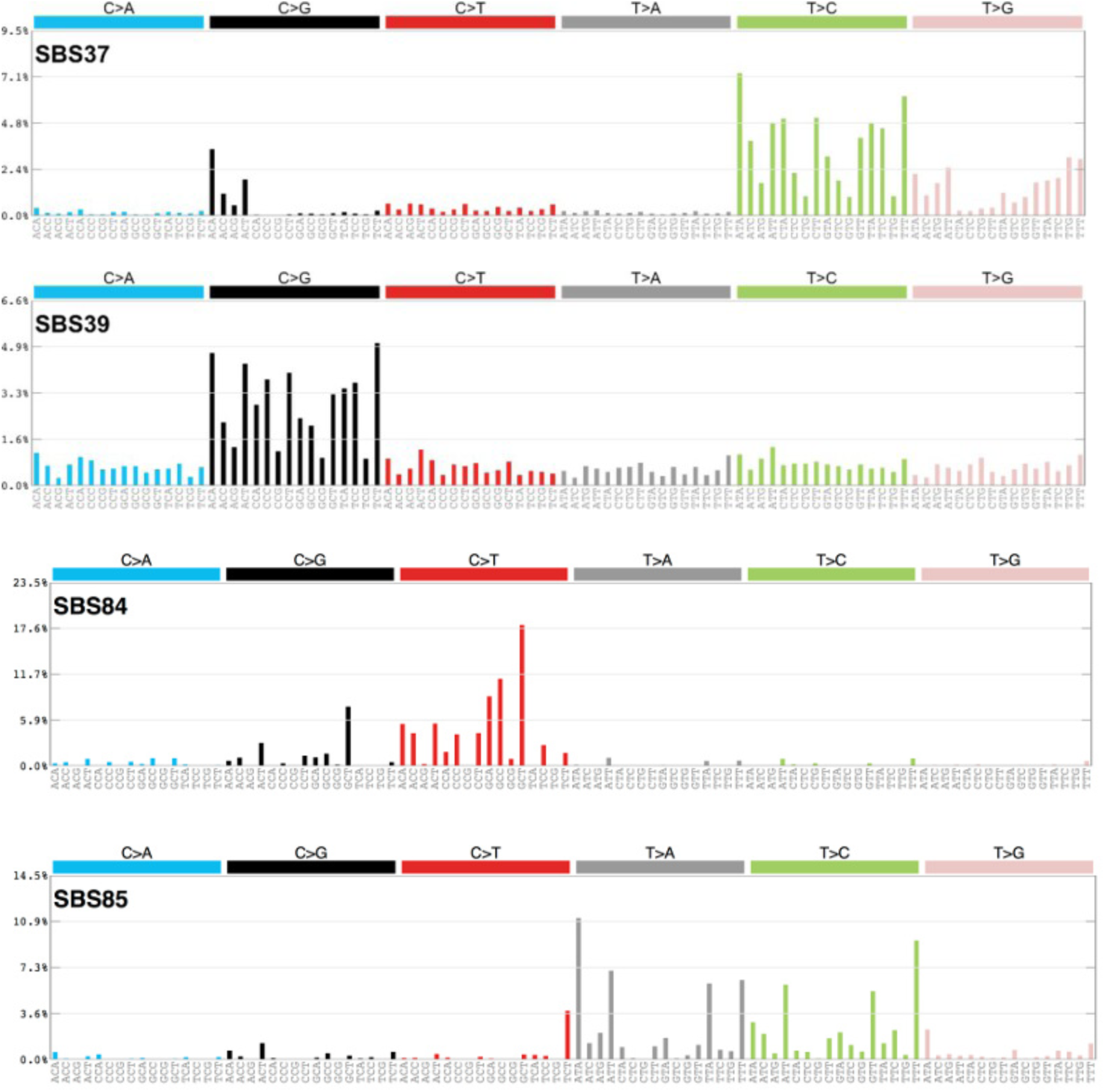
COSMIC SBS signatures related to SHM. SBS 84 and SBS 85 are denoted as signatures of AID-induced somatic hypermutations in lymphoid cells, whereas the etiology of SBS 37 and SBS 39 is listed as unknown in COSMICv3. Our data suggest that SBS 37 and SBS 39 are likely signatures of error-prone DNA polymerases in SHM. These graphs are adapted from the COSMIC website for reference.

**Figure S11.**
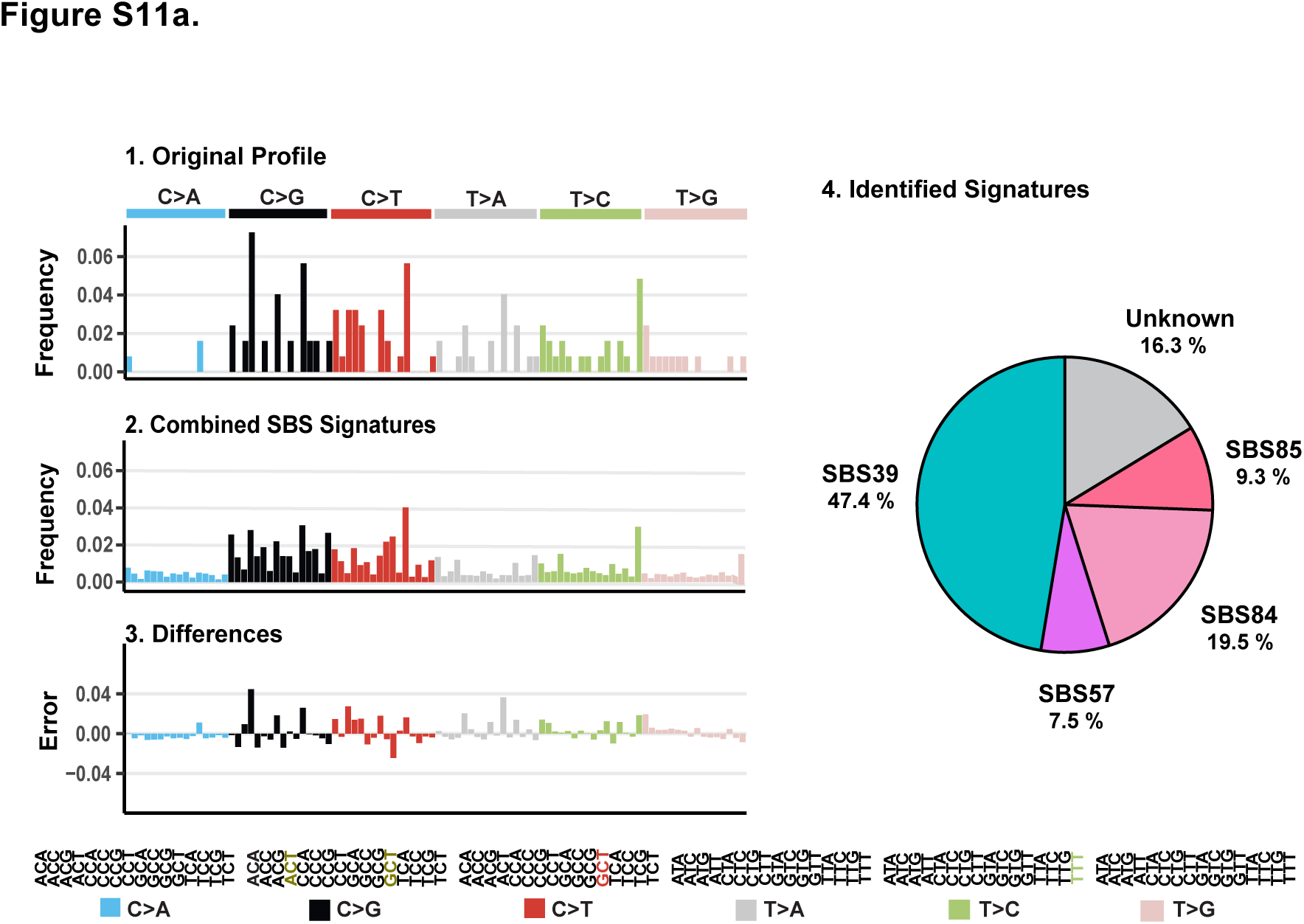
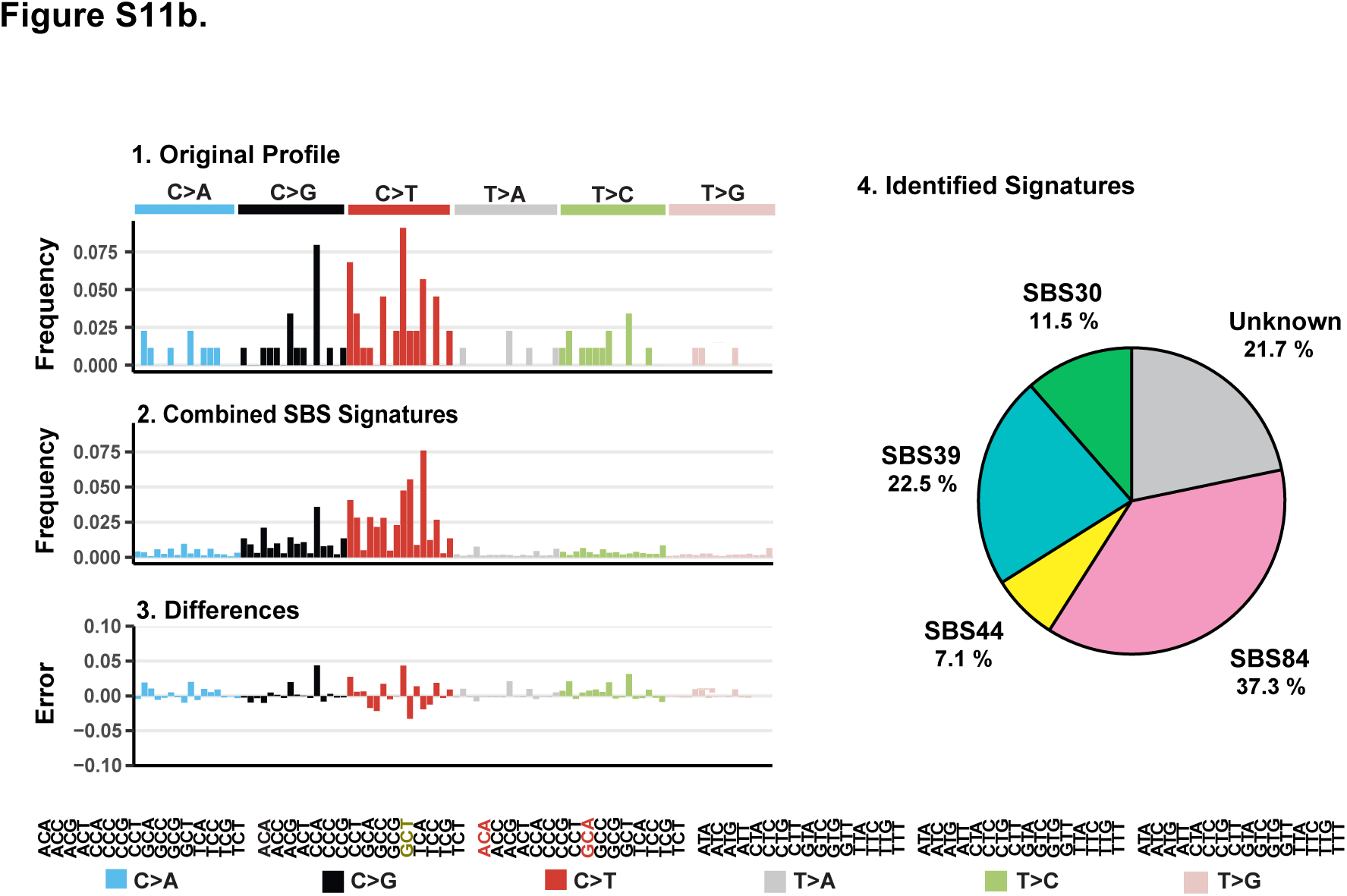
Mutational signatures of *PAX5* SE (a) and *CD83* SE (b). Panels represent the original profiles (a-1 and b-1), the signature as computed using the best fit combination of COSMICv3 SBS signatures (a-2 and b-2), the difference between the preceding two figures (a-3 and b-3), and the breakdown of the original profile into COSMIC v3 SBS signatures (a-4, b-4). Types of base substitutions and the array of trinucleotide motifs are displayed; motifs with the most frequent substitutions found in the original profile (panel 1) are highlighted with colors designated for each of the six base substitution types.

### Supplementary Tables

**Table S1.** Gene loci used for the design of DNA capture probes.

|  | Gene | Chromosome | Position |
| --- | --- | --- | --- |
| 1 | *BCL6* | chr3 | 187,462,513 - 187,463,513 |
| 2 | *RHOH* | chr4 | 40,192,631 - 40,193,958 |
| 3 | *EBF1* | chr5 | 158,525,438 - 158,526,788 |
| 4 | *CD83* | chr6 | 14,117,487 - 14,118,296 |
| 5 | *PIM1* | chr6 | 37,138,549 - 37,139,267 |
| 6 | *MYC* | chr8 | 128,748,315 - 128,749,694 |
| 7 | *PAX5* | chr9 | 37,033,467 - 37,034,476 |
| 8 | *POU2AF* | chr11 | 111,248,887 - 111,250,157 |
| 9 | *H2AFX* | chr11 | 118,964,585 - 118,966,177 |
| 10 | *AICDA* | chr12 | 8,764,442 - 8,765,442 |
| 11 | *CD79B* | chr17 | 62,008,695 - 62,009,704 |
| 12 | *CD79A* | chr19 | 42,381,190 - 42,382,190 |
| 13 | *IgHV3-23* | chr14 | 106,725,101 - 106,725,833 |

**Table S2.** Donor information.

| **Individual** | **Age** | **Race** | **Smoker** |
| --- | --- | --- | --- |
| **1** | **32** | **Caucasian** | **No Information** |
| **2** | **30** | **Hispanic** | **No Information** |
| **3** | **29** | **Asian** | **No** |
| **4** | **24** | **African American** | **No** |
| **5** | **53** | **African American** | **No** |
| **6** | **22** | **Asian** | **No** |
| **7** | **24** | **Caucasian** | **No** |
| **8** | **25** | **Hispanic** | **No** |
| **9** | **26** | **Caucasian** | **No** |
| **10** | **24** | **Hispanic** | **No** |

**Table S3.** Average mutation frequencies in various super-enhancer loci in CD19+ B-cells from ten healthy individuals. The SEs with the highest and the lowest mutation frequency are highlighted.

| Gene | Total Mutations Detected | Total Nucleotides Read | Mutation Frequency |
| --- | --- | --- | --- |
| *BCL6* | 7,853 | 35,565,417 | $2.2 \times 10^{-4}$ |
| *RHOH* | 12 | 23,515,071 | $5.1 \times 10^{-7}$ |
| *EBF1* | 14 | 19,173,584 | $7.3 \times 10^{-7}$ |
| *CD83* | 198 | 28,562,424 | $6.9 \times 10^{-6}$ |
| *PIM1* | 68 | 26,048,381 | $2.6 \times 10^{-6}$ |
| *MYC* | 64 | 47,415,092 | $1.3 \times 10^{-6}$ |
| *PAX5* | 259 | 26,690,301 | $9.7 \times 10^{-6}$ |
| *POU2AF* | 405 | 43,216,310 | $9.4 \times 10^{-6}$ |
| *H2AFX* | 13 | 45,842,880 | $2.8 \times 10^{-7}$ |
| *AICDA* | 29 | 33,710,036 | $8.6 \times 10^{-7}$ |
| *CD79B* | 83 | 39,229,867 | $2.1 \times 10^{-6}$ |
| *CD79A* | 15 | 25,015,537 | $6.0 \times 10^{-7}$ |
| *IgHV3-23* | 2,059 | 16,423,011 | $1.3 \times 10^{-4}$ |

**Table S4.** Average mutation frequencies of *BCL6* SE in different ethnic groups.

| **Ethnic Group** | **Individual** | **Age** | **Mutation Frequency**<br>**(*BCL6* SE)** | **Average**<br>**Mutation Frequency** |
| --- | --- | --- | --- | --- |
| **Caucasian** | **1** | **32** | **$1.2 \times 10^{-4}$** | **$1.5 \times 10^{-4}$** |
|  | **7** | **24** | **$2.6 \times 10^{-4}$** |  |
|  | **9** | **26** | **$6.4 \times 10^{-5}$** |  |
| **Hispanic** | **2** | **30** | **$2.5 \times 10^{-4}$** | **$2.0 \times 10^{-4}$** |
|  | **8** | **25** | **$2.5 \times 10^{-4}$** |  |
|  | **10** | **24** | **$1.1 \times 10^{-4}$** |  |
| **African American** | **4** | **24** | **$2.2 \times 10^{-4}$** | **$4.3 \times 10^{-4}$** |
|  | **5** | **53** | **$6.3 \times 10^{-4}$** |  |
| **Asian** | **3** | **29** | **$2.1 \times 10^{-4}$** | **$2.0 \times 10^{-4}$** |
|  | **6** | **22** | **$1.9 \times 10^{-4}$** |  |

## References

1. Klein U, Dalla-Favera R (2008) Germinal centres: role in B-cell physiology and malignancy. Nat Rev Immunol 8(1):22–33.

2. Victora GD, Nussenzweig MC (2012) Germinal Centers. Ann Rev Immunol 30:429–57.

3. De Silva NS, Klein U (2015) Dynamics of B cells in germinal centres. Nat Rev Immunol 15(3):137–148.

4. Peled JU, et al. (2008) The biochemistry of somatic hypermutation. Ann Rev Immunol 26:481–511.

5. Liu MM., et al. (2008) Two levels of protection for the B cell genome during somatic hypermutation. Nature 451(7180):841–845.

6. Meng FL, et al. (2014) Convergent transcription at intragenic super-enhancers targets AID-initiated genomic instability. Cell 159(7):1538–1548.

7. Qian J, et al. (2014) B cell super-enhancers and regulatory clusters recruit AID tumorigenic activity. Cell 159(7):1524–1537.

8. Pasqualucci L, et al. (1998) BCL-6 mutations in normal germinal center B cells: evidence of somatic hypermutation acting outside Ig loci. Proc Natl Acad Sci U S A 95(20):11816–11821.

9. Shen HM, Peters A, Baron B, Zhu X, Storb U (1998) Mutation of BCL-6 gene in normal B cells by the process of somatic hypermutation of Ig genes. Science 280(5370):1750–1752.

10. Crotty S, Johnston RJ, Schoenberger SP (2010) Effectors and memories: Bcl-6 and Blimp-1 in T and B lymphocyte differentiation. Nat Immunol 11(2):114–120.

11. Basso K, Dalla-Favera R (2012) Roles of BCL6 in normal and transformed germinal center B cells. Immunol Rev 247(1):172–183.

12. Migliazza A, et al. (1995) Frequent somatic hypermutation of the 5’ noncoding region of the BCL6 gene in B-cell lymphoma. Proc Natl Acad Sci U S A 92(26):12520–12524.

13. Basso K, Dalla-Favera R (2015) Germinal centres and B cell lymphomagenesis. Nat Rev Immunol 15(3):172–84.

14. Saito M, et al. (2007) A Signaling Pathway Mediating Downregulation of BCL6 in Germinal Center B Cells Is Blocked by BCL6 Gene Alterations in B Cell Lymphoma. Cancer Cell 12(3):280–292.

15. Schmitt MW, et al. (2012) Detection of ultra-rare mutations by next-generation sequencing. Proc Natl Acad Sci U S A 109(36):14508–14513.

16. Kennedy SR, et al. (2014) Detecting ultralow-frequency mutations by Duplex Sequencing. Nat Protoc 9:2586–2606.

17. Wu YC, et al. (2010) High-throughput immunoglobulin repertoire analysis distinguishes between human IgM memory and switched memory B-cell populations. Blood 116(7):1070–1078.

18. Weller S, et al. (2001) CD40-CD40L independent Ig gene hypermutation suggests a second B cell diversification pathway in humans. Proc Natl Acad Sci U S A 98(3):1166–1170.

19. Morbach H, Eichhorn EM, Liese JG, Girschick HJ (2010) Reference values for B cell subpopulations from infancy to adulthood. Clin Exp Immunol 162(2):271–279.

20. Lu Z, et al. (2013) BCL6 breaks occur at different AID sequence motifs in Ig-BCL6 and non-Ig-BCL6 rearrangements. Blood 121(22):4551–4554.

21. Lu Z, et al. (2015) Convergent BCL6 and lncRNA promoters demarcate the major breakpoint region for BCL6 translocations. Blood 126(14):1730–1731.

22. Matsuda T, Bebenek K, Masultanl C, Hanaoka F, Kunkel TA (2000) Low fidelity DNA synthesis by human DNA polymerase-η. Nature 404(6781):1011–1013.

23. Loeb LA, Preston BD (2003) Mutagenesis by Apurinic/Apyrimidinic Sites. Ann Rev Genet 20(1):201–230.

24. Kano C, Hanaoka F, Wang JY (2012) Analysis of mice deficient in both REV1 catalytic activity and POLH reveals an unexpected role for POLH in the generation of C to g and G to C transversions during Ig gene hypermutation. Int Immunol 24(3):169–174.

25. Masuda K, et al. (2009) A Critical Role for REV1 in Regulating the Induction of C:G Transitions and A:T Mutations during Ig Gene Hypermutation. J Immunol 183:1846–1850.

26. Alexandrov LB, et al. (2013) Signatures of mutational processes in human cancer. Nature 500(7463):415–21.

27. Rosenthal R, McGranahan N, Herrero J, Taylor BS, Swanton C (2016) deconstructSigs: Delineating mutational processes in single tumors distinguishes DNA repair deficiencies and patterns of carcinoma evolution. Genome Biol. doi:10.1186/s13059-016-0893-4.

28. Zhang L, et al. (2019) Single-cell whole-genome sequencing reveals the functional landscape of somatic mutations in B lymphocytes across the human lifespan. Proc Natl Acad Sci U S A 116(18):9014–9019.

29. Rogozin IB, Kolchanov NA (1992) Somatic hypermutagenesis in immunoglobulin genes. BBA - Gene Struct Expr 1171(1):11–18.

30. Zhao Y, et al. (2013) Mechanism of somatic hypermutation at the WA motif by human DNA polymerase η. Proc Natl Acad Sci U S A 110(20):8146–51.

31. Zeng X, et al. (2001) DNA polymerase eta is an A-T mutator in somatic hypermutation of immunoglobulin variable genes. Nat Immunol 2(6):537–541.

32. Rogozin IB, Pavlov YI, Bebenek K, Matsuda T, Kunkel T a (2001) Somatic mutation hotspots correlate with DNA polymerase eta error spectrum. Nat Immunol 2(6):530–536.

33. Seki M, Gearhart PJ, Wood RD (2005) DNA polymerases and somatic hypermutation of immunoglobulin genes. EMBO Rep 6(12):1143–1148.

34. Zanotti KJ, Gearhart PJ (2016) Antibody diversification caused by disrupted mismatch repair and promiscuous DNA polymerases. DNA Repair 38:110–116.

35. Storb U, Stavnezer J (2002) Immunoglobulin genes: Generating diversity with AID and UNG. Curr Biol 12(21):R725–R727.

36. Keim C, Kazadi D, Rothschild G, Basu U (2013) Regulation of AID, the B-cell genome mutator. Gene Dev 27(1):1–17.

37. Zan H, et al. (2001) The translesion DNA polymerase ζ plays a major role in Ig and bcl-6 somatic hypermutation. Immunity 14(5):643–653.

38. Zan H, et al. (2005) The translesion DNA polymerase θ plays a dominant role in immunoglobulin gene somatic hypermutation. EMBO J 24(21):3757–3769.

39. Weill J-C, Reynaud C-A (2008) DNA polymerases in adaptive immunity. Nat Rev Immunol 8(4):302–312.

40. Ouchida R, et al. (2008) Genetic analysis reveals an intrinsic property of the germinal center B cells to generate A:T mutations. DNA Repair 7(8):1392–1398.

41. Masuda K, et al. (2008) DNA polymerase η is a limiting factor for A:T mutations in Ig genes and contributes to antibody affinity maturation. Eur J Immunol 38(10):2796–2805.

42. Mcheyzer-Williams LJ, Milpied PJ, Okitsu SL, Mcheyzer-Williams MG (2015) Class-switched memory B cells remodel BCRs within secondary germinal centers. Nat Immunol 16:296–305.

43. Supek F, Lehner B (2017) Clustered Mutation Signatures Reveal that Error-Prone DNA Repair Targets Mutations to Active Genes. Cell 170:534–547.

44. Eid MMA, et al. (2017) Integrity of immunoglobulin variable regions is supported by GANP during AID-induced somatic hypermutation in germinal center B cells. Int Immunol 29(5):211–220.

45. Calo E, Wysocka J (2013) Modification of Enhancer Chromatin: What, How, and Why? Mol Cell 49(5):825–837.

46. Lieber MR (2016) Mechanisms of human lymphoid chromosomal translocations. Nat Rev Cancer 16(6):387–398.

47. Ye BH, et al. (1995) Chromosomal translocations cause deregulated BCL6 expression by promoter substitution in B cell lymphoma. EMBO J 14(24):6209– 6217.

48. Hatzi K, Melnick A (2014) Breaking bad in the germinal center: How deregulation of BCL6 contributes to lymphomagenesis. Trends Mol Med 20(6):343–352.

49. Martincorena I, et al. (2018) Somatic mutant clones colonize the human esophagus with age. Science 362(6417):911–917.

50. Yokoyama A, et al. (2019) Age-related remodelling of oesophageal epithelia by mutated cancer drivers. Nature 565(7739):312–317.

51. Yizhak K, et al. (2019) RNA sequence analysis reveals macroscopic somatic clonal expansion across normal tissues. Science 364(6444):doi: 10.1126/science.aaw0726.

52. Schmitt MW, et al. (2015) Sequencing small genomic targets with high efficiency and extreme accuracy. Nat Methods 12(5):423–425.

